# Challenges of γ9δ2TCR affinity maturation when using phage display

**DOI:** 10.1101/2021.05.20.445024

**Authors:** Lovro Kramer, Margaux Demuysere, Eline van Diest, Dennis X. Beringer, Jürgen Kuball

## Abstract

**Background:** Over the past years we showed that the efficacy of αβT cells engineered to express a defined γδTCR (TEG) depends on the functional avidity of the γ9δ2TCR. We hypothesized that functional avidity mediated through γ9δ2TCR in the TEG format could be further enhanced by increasing affinity of the γ9δ2TCR.

**Methods:** We attempted to overcome limited affinity of natural occurring γ9δ2TCRs through affinity maturation by phage display using a library containing mutations in CDR1 and CDR2 of both TCR chains.

**Conclusion:** Affinity maturation of γ9δ2TCR by using phage display was not successful. The largest hurdle was the periplasmic expression of γ9δ2TCR constructs in *E.coli* which is a prerequisite for successful phage display. The underlying reason for this lack of expression was the instability of the single chain (sc)TCR format. Expression of scTCR formats in HEK293F cells yielded only 15-20% correctly folded scTCR.

## Introduction

Over the past years we showed that the efficacy of αβT cells engineered to express a defined γ9δ2TCR (TEG) is modulated through the functional avidity of the γ9δ2TCR (1,2). Functional avidity consists of the affinity the TCR for its ligand as well as the absolute number of receptors on the cell surface. We showed that by increasing the surface expression of the γ9δ2TCR functional avidity of TEGs increased (3). Increasing affinity of the γ9δ2TCR would be the next logical step to increase functional avidity of TEGs. Increasing affinity of the γ9δ2TCR has however the risk of inducing auto-reactivity as the ligands identified so far seem to be ubiquitously expressed in all tissues (1). Regardless of this therapeutic risk, γ9δ2TCR with increased affinity would be useful as improved staining tools to identify cells to be potentially targeted by TEGs. To date, γ9δ2TCRs in a tetramer or bead format, do not recognize all tumor cells targeted by TEGs (1).

One approach to tackle limited affinity of natural occurring TCRs is to enhance their affinity by mutagenesis. For many αβTCRs this strategy has been used successfully and several of these αβTCRs entered clinical testing (4,5). A recent clinical trial using αβT cells engineered with high-affinity NY-ESO αβTCR has demonstrated a clinical benefit superior to the wild-type TCR, highlighting the power of affinity maturation (5). The great strategic advantage from a designer perspective for affinity maturation of αβTCRs is that the ligands of an αβTCR are well defined. In contrast, even with the recent advances in understanding key molecules and pathways for the activation of γ9δ2T cells, there is no clearly defined ligand-complex for γ9δ2TCRs (6–10). Within this context we describe challenges when using a construct comprising covalently linked, single chain, γ9δ2 variable domains (scTCR) for the affinity maturation through phage display.

## Methods

### Cloning, Expression and purification of single chain Clone 5 in HEK293F cells

Synthetic DNA for a scTCR construct of Clone 5 (scCl5) was synthesized by BaseClear including 5 mutations in TCRγ9 and 1 mutation in TCRδ2 based on important stabilizing mutations in scαβTCRs (11) (See appendix 1 Sequences: “scCl5-HEK”). The synthetic gene was sub-cloned, using BswI and SalI restriction sites, in a modified pcDNA3 vector (kind gift from protein facility LTI; UMCU), containing a 3’ biotin acceptor peptide and His-tag after the SalI resitriction site.

HEK293 freestyle cells were transfected with this expression construct in combination with pBu-BirA using 293Fectin (Thermo fisher scientific) transfection reagent at 1μg DNA/10^6 cells. After 6 hours the media was supplemented with 0.5 % Pen/Strep and 100 μM D-biotin and the transfected cells were cultured for 5 days, after which the expression media was harvested.

The scCl5 was purified using a two-steps protocol using first immobilized metal affinity chromatography (IMAC) to enrich for His tagged protein (1 ml HisTrap, GE Healthcare) followed by anion exchange (1 ml HiTrapQ, GE Healthcare), according to manufacturers instructions. Collected fractions were analyzed by SDS-PAGE and pooled and concentrated based on the chromatogram and SDS-PAGE analysis.

### Staining of cell lines using scCl5-beads

Biotinylated soluble TCR Cl5 (1) or scCl5 were mixed with 5-7μm streptavidin-coated UV-beads (Spherotech) in excess to ensure fully coated beads, 10 μg sTCR/mg microspheres.

7.5*104 cells, ML1 or K562, were incubated with 20 μl Cl5-UV beads (0.33 mg beads/ml) for 30 minutes at RT. The mixtures were fixed by adding 20 μl 2% formaldehyde for 15 minutes. Samples were washed once with 1% formaldehyde and analyzed on a BD FACSCanto II (BD).

### Expression of scCl5 in E.coli JM109 using phagemid pUR8100

*E.coli* codon optimized DNA encoding scCl5 (Vδ2-linker Vγ9) was ordered at BaseClear and sub-cloned in to phagemid pUR8100 (kind gift QVQ/Mohamed El Khattabi) using SfiI and NotI restriction sites (See appendix 1 Sequences: “scCl5-Ecoli”).

Chemical competent *E.coli* JM109 were transformed with pUR8100-scCl5 and selected on 2YT-agar plates supplemented with 100 μg/ml ampicillin and 0.1% v/v glucose. A single colony was used to inoculate 50 ml 2YT (+ ampicillin & 0.1% v/v glucose) for overnight growth at 37°C. 5 ml of the overnight culture was used to inoculate 1 l 2YT (+ ampicillin), once the culture reached an OD_600nm_ of 0.6, isopropyl β-D-1-thiogalactopyranoside (IPTG) was added to a final concentration of 0.5 mM. Three hours after IPTG induction the bacteria were pelleted and the dry pellets were frozen at −20°C overnight. The cells were thawed and resuspended 100 ml in IMAC buffer and incubated for 30’ on ice. The cells were pelleted (13,000x*g* for 20’) and the supernatant was filtered over a 0.2 μm filter. Further purification was done as described above. The anion exchange fractions were analyzed by Western blot using anti-His antibody (BD Pharmingen; Cat#552565) and Goat poyclonal anti-mouse IgG-HRP (Santa Cruz Biotechnology; sc2005).

### Assembly of the scCl5 D12G12 library

For the assembly of the scCl5 D12G12 library several consecutive PCRs had to be done to introduce the mutagenic primers. Primer sequences used are listed in Table S1 (See appendix 1 Sequences).

First four megaprimers were amplified from using scCl5-Ecoli as template: “megaprimer 1a” was amplified using primers pMP1_f_2 and pMP1_r, “megaprimer 1b” was amplified using primers pMP2_f and pMP2_r, “megaprimer 1c” was amplified using primers pMP3_f and pMP3_r_2, and “megaprimer 1d” was amplified using primers pMP4_f and pMP4_r. Phusion DNA polymerase was used to amplify the megaprimers using a standard protocol with an annealing temperature of 52°C and 35 cycles were done.

These megaprimers were used as templates for PCR to introduce the mutagenic primers using the following concentrations of primers and template: upstream primer 0.5 μM, downstream primer 0.5 μM, mutagenic primer 25 nM, and megaprimer 25 nM. Again Phusion DNA polymerase was used, for these reactions the annealing temperature was 60°C and 35 cycles were done. The sublibrary for CDR1δ was constructed using pMP1_f_2, pdCDR1-linkd12_r2, pdCDR1mut, and megaprimer 1a. The sublibrary for CDR2δ was constructed using plinkd12-dCDR2_f2, pdCDR2-Kranz_r, pdCDR2mut, and megaprimer 1b. The sublibrary for CDR1γ was constructed using pKranz-gCDR1_2, pgCDR1-linkg12_r2, pgCDR1mut, and megaprimer 1c. The sublibrary for CDR2γ was constructed using plinkg12-gCDR2_f, pCtermNotI_2, pgCDR2mut, and megaprimer 1d. Products CDR1δ and CDR1γ were digested with BsaI-HF, and CDR2δ and CDR2γ with BbsI-HF. CDR1δ and CDR2δ were mixed at equimolar ratio and ligated using T4 ligase, same was done for CDR1γ and CDR2γ. The ligated products CDR1δ+CDR2δ “D12” and CDR1γ+CDR2γ “G12” were amplified using PCR with phusion DNA polymerase using primers pMP1_f_2 and pdCDR2-Kranz_r for “D12” and primers pKranz-gCDR1_2 and pCtermNotI_2 for “G12”. The purified PCR products were digested with BsaI-HF for “D12”, and with BbsI-HF for “G12”. The two digested products were ligated with T4 ligase and amplified by PCR using primers pNtermSfi_2 and pCtermNotI_2 to create the final library scCl5“D12G12”.

The scCl5“D12G12” PCR product was digested with SfiI and NotI and ligated in pUR8100 that was also digested with SfiI and NotI, and dephosphorylated with rSAP. 315 ng scCl5“D12G12” and 450 ng pUR8100 was mixed and incubated with T4 ligase for 3 hours at room temperature, the ligase was heat inactivated at 65°C for 10 minutes.

The ligated pUR8100-scCl5”D12G12” was used to transform *E.coli* XL1-blue, to assess ligation efficiency and library quality. Phagemids of 6 individual colonies were analyzed by sanger sequencing.

### Preparation of scCl5”D12G12”-displaying bacteriophages

On the first day, the strain *E.coli* XL-1 blue was transformed via heat-shock with the phagemids containing the scCl5”D12G12”. The transformed bacteria were grown o/n in 5mL of 2xYT medium supplemented with Amp100 and Glc1%. On day 2, the o/n growth culture was diluted into fresh 2xYT medium supplemented with Amp100 and Glc1% and grown to an OD_600nm_ of 0.5. The bacteria were then infected with 2*10^11 plaque forming units (pfu) of M13KO7 helper phage and grown o/n at 37°C in fresh 2xYT medium supplemented with Amp100 and 75μg/mL of kanamycin (Kan75, Gibco) to select for bacteria containing a phagemid and infected with a helper phage. The medium was devoid of glucose to enable the expression of the scCl5”D12G12”-gIII fusion protein and thus the encapsidation of helper phages displaying the scCl5”D12G12”. On day 3, the recombinant phages particles were precipitated twice with a solution of PEG 8000 20% (Fisher)/NaCl 2.5M and pelleted each time by a centrifugation step of 10 minutes at 4°C/3200xg to remove all bacterial contaminants. 20% glycerol (Sigma Aldrich) stocks were made for further use, the remaining phages were stored in at 4°C up to one week.

### Phage titer determination

*E.coli* TG1 were streaked on a 2xYT agar plate supplemented and Glc1%. One colony was grown in 2xYT medium supplemented with Glc1% at 37°C/200rpm until the OD_600nm_ reached 0.4 to 0.5. A serial log-dilution of phage particles in PBS was added (1:10) to the bacterial culture dived over different tubes. 5μL of the infected bacteria at different dilutions and PBS only as a negative control were spotted on a 2xYT agar plate supplemented with Amp100 and Glc1% and incubated o/n at 37°C.

The phage titer (pfu/mL) was then determined as the mean for different log-dilutions of:

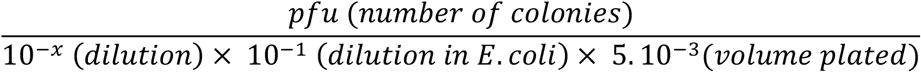

### Bio-panning of scCl5”D12G12” library

5×10e9 pfu of the phage library was diluted 1:1 with PBS-2% milk and incubated for 15 min at RT. 3*10^6 ML1 cells were washed with PBS and resuspended in 1 mL PBS-2% milk and the diluted phages were added and incubate for 30min at RT and slow rotation. The unbound phage fraction was collected by centrifugation at 400g and careful aspiration of the supernatant. K562 cells, preincubated with 100 uM pamidonate, were washed with PBS and resuspended them in 1 mL PBS-milk2%. The unbound phage fraction was added and incubated for 30 min at RT and slow rotation (panning). The cells 4 times with PBS-2%milk (centrifugation for 5min at 400g) followed by 4 additional wash steps with PBS. Bound phages were eluted by resuspending the cells in 1 mL of 0.1M Glycine/HCl (pH 2.2). The tube was rotated for 10 min on RT and centrifuged at 4000g for 2 min. Immediately, the supernatant was collected and neutralized with 100 uL of 2M TRIS/HCl (pH 8.0). The eluted and neutralized phages were transferred to a microcentrifuge tube and centrifuged at 20000g for 5 min at RT. Collect the supernatant. 20 μL of the eluted phages was used for analysis of phage titer. The remaining phage solution was used to infect *E. coli* cells and for the preparation of glycerol stocks.

### Binding analysis of single scCl5”D12G12” clones with flow cytometry

94 single colonies of *E.coli* infected with K562-eluted phages were transferred to a 96-well plate containing 1ml/well of 2xYT medium supplemented with Amp100 and Glc1% and grown overnight. The following day two new plates were inoculated with 100 μl overnight culture. scCl5”D12G12”Myc-His was induced at OD_600nm_ of 0.6-0.8 with 0.5 mM IPTG and scCl5”D12G12” phages were produced by infection with M13KO7 helper phage. scCl5”D12G12” and scCl5”D12G12”-phages were prepared as described above and used to incubate ML1 and K562 cells. Following washing, the binding of scCl5”D12G12” and scCl5”D12G12”phage was assessed via incubation at RT with an anti-Myc or an anti-M13 antibodies respectively. The antibodies were diluted to a final concentration of 1.7ng/mL in PBS. Both antibodies are biotinylated and were therefore detected with a streptavidin-phycoertyhrin (SA-PE) conjugate solution (diluted 1:1000 in a 1% w/v BSA fraction V (Sigma-Aldrich)/PBS solution). The viability of the cells was also assessed via eBioscienceTM Fixable Viability Dye eFluorTM 780. The stained cells were fixed with BD CytofixTM Fixation Buffer and measured with FACSCanto II. The measurements were analyzed on Diva Software.

### Construction of bacterial expression plasmids for double-chain γδTRCs

The sequences from the γ9δ2TCR Cl5 and the γ4δ5TCR C132 were optimized for bacterial expression and modified according to the double chain (dc)TCR design invented by Jakobsen and Glick (12). Briefly, the ectodomains from each chain were conserved up to the amino acids preceding the membrane-proximal disulfide bond. The amino acids glutamine 79 and valine 84 from the γ and δ constant domains respectively were replaced by a cysteine, thereby introducing a non-native disulfide bond to stabilize the interaction between both chains. A Biotin Acceptor Peptide (BAP) tag was annexed C-terminally to the δ chain of dcCL5. A periplasmic leader sequence, either gene III (gIII) or pectate lyase B (pelB), was added to the N-terminus of the γ-chain of CL5 and C132 respectively while a pelB sequence was already present into the host plasmids in-frame with the δ-chain. Each genetic construct was flanked with the restriction sites SfiI and NotI (See appendix 1 Sequences: “dcCl5-Ecoli” and “dcC132-Ecoli”).

Both constructs were subcloned into the phagemid pUR8100 and its derivative, the expression vector pMEK219 that is devoid of the the pIII protein. Both vectors drive the expression of the dcTCR from the lac promoter. They additionally contain two C-terminus tags (myc and his) in-frame with the γ-chain that can be subsequently used for detection and/or purification.

### Small scale expression test of dcTCRs in E.coli XL1-blue

Transformed bacteria were grown overnight (o/n) in 5 mL of 2x YT (tryptone + yeast extract + NaCl BioXtra, Sigma Aldrich) medium supplemented with Amp100 and 1% of glucose (Glc1%) in a shaking incubator at 37°C/200rpm. On the next morning, 500μL of the o/n growth culture was inoculated to 50mL of fresh 2x YT medium supplemented with Amp100 and grown at 37°C/200rpm to an OD600nm of 0.6-0.8. The expression of the dcTCR was then induced via the addition of 0.1mM-1M of IPTG to the culture medium followed by an incubation either of 3h at 37°C/200rpm or o/n at 20°C or 30°C/200rpm. Large-scale expression was performed essentially as detailed above. The o/n culture volumes were upscaled to 200mL from which 2mL were inoculated into 200mL of fresh 2xYT medium for protein expression. The expression was induced with the addition of IPTG to a final concentration of 0.5mM and expressed o/n at 20°C/200rpm.

The samples for cytosolic and periplasmic extraction were isolated in parallel before and after protein induction. Upon collection, they were centrifuged at 6000xg for 10 minutes at room temperature (RT). The supernatant was carefully discarded and the bacterial pellets were frozen o/n. On the following day, the samples were thawed at RT for 15 minutes. The periplasmic extracts were prepared by resuspending the cell pellets in the extraction buffer (20mM Tris/HCl, 300mM NaCl, pH 8.2 supplemented with proteases inhibitor. The resuspended samples were incubated on ice while shaking for 30 minutes, before a final centrifugation step at 4°C/13,000xg for 20 minutes. The supernatant was collected and further analyzed by Western blot as described above.

### Generation of randomly mutated scTCR

scCl5 and scC132 were cloned in the periplasmic expression vector pMEK219 using SfiI and NotI restriction sites (See appendix 1 Sequences: “scCl5-Ecoli” and “scC132-Ecoli”). These plasmids were used as template for random mutagenesis. The protocol used to introduce mutations within the scTCR sequence was adapted from a previously published report aiming to generate scTCR mutants for yeast display (13). Briefly, the wild-type scTCR construct was amplified from 50ng of pMEK219-scTCR plasmid (the primers used can be found in the Supplemental Table 2) with a Taq polymerase (Invitrogen) in presence of a dNTP imbalance (220 μM of dATP, 200 μM of dCTP, 340 μM of dGTP and 2.4 mM of dTTP), an excess of magnesium (Magnesium chloride, Invitrogen) and the mutagenic compound manganese (Manganese chloride solution, Sigma Aldrich). 10μL of the reaction were run on gel to confirm the gene amplification. The PCR product was digested with DpnI to eliminate the template DNA, then purified and concentrated with the NucleoSpin^®^ Gel and PCR Clean-up kit (Macherey Nagel). Subsequently, the mutated PCR fragment was reintroduced into the desired plasmid using the aforementioned restriction sites.

### Expression and periplasmic extraction scCl5, scC132 and nanobody

Colonies of the E.coli strains TOP10 and DH5-α transformed with the wild-type genes (Nb, scCL5, scC132) or the library of mutants respectively were picked and transferred in a 96-deep well plate (Eppendorf) containing 1mL of LB medium (MP Biomedicals) supplemented with Amp100. The plate was left shaking o/n in the incubator at 37°C/200rpm. On the next day, the OD600nm of some clones was measured via a spectrophotometer (Eppendorf) and the cultures were diluted into fresh LB-Amp100 medium to adjust the OD600nm around 0.1. The new 96-well plate was grown at 37°C/200rpm to an OD600nm of 0.6-0.8 while the remaining culture was kept in the fridge for further sequencing. The expression of the different proteins was then induced via the addition of 0.5mM of IPTG to each well followed by an incubation of 3h at 37°C. The OD600nm of different wells was assessed again. The 96-deep well plate was spun down for 5 minutes at RT/3200rpm. The supernatant was carefully removed and the sealed plate was stored in the freezer. In the following days, the plate was thawed at RT for 15 minutes. The cell pellets were resuspended in 100μL of PBS and transferred to a 96-well plate with V-bottom tubes (Greiner bio-one). The V-bottom plate was centrifuged for 10 minutes at RT/3200rpm. The supernatants were kept on ice for 30 minutes before being further analyzed as explained below.

### Analysis of scTCR expression by dot-blot

The supernatants were collected and analyzed by means of a Bio-Dot apparatus (Bio-Rad) according to the manufacturer’s instructions. Briefly, the proteins were directly blotted against a pre-wet nitrocellulose membrane using both gravitational and aspiration forces. The membrane was blocked for 1 hour in PBS supplemented with 1% Tween-20 (Merck MP Biomedicals) and 5% skimmed milk (Friesland Campina). The dcTCR were detected with a primary antibody anti-his (1:1000 dilution) and a secondary antibody antimouse (1:5000 dilution) conjugated to HRP in PBS supplemented with 1% Tween-20 and 1% skimmed milk. The membrane was washed and developed with Chemiluminescent Peroxidase Substrate-1 (Sigma-Aldrich).

## Results

### A fraction of single chain γ9δ2TCR expressed in HEK293F can bind K562 as soluble γ9δ2TCR

Single chain variable domains are the most commonly used antibody and/or TCR fragments used to be incorporated in phage coat proteins for display purposes. A single chain γ9δ2TCR construct of clone 5, containing the Vδ2 domain linked to the Vγ9 domain linked to a biotin acceptor peptide and a His-tag (figure 1A) was transiently expressed in HEK293F cells. Six days after transfection single chain Cl5, scCl5, was purified using a nickel column (his-tag purification) followed by an anion exchange column (charge-based separation). The chromatogram of the anion exchange showed many different peaks (figure 1B), indicating the presence of several scCl5 charge variants. Charge differences of expressed proteins can have one or more causes, like differential glycosylation, other post translational modifications, degradation, and/or incorrect folding. The different fractions were analyzed by SDS-PAGE to determine if there was a obvious cause of the many different charge variants. The majority of the fractions showed an intense band around 30 kDa, which is the expected size of scCl5 (figure 1C). Fractions 11-13 also contained a less intense band around 60 kDa, which might indicate the presence of covalently linked dimers, e.g. via a disulfide bond, and fraction 27 contained a broad smear between 37 and 50 kDa. Apart from the last peak, containing fraction 27, all other peaks contained protein of similar size, sometimes with minor contaminations. To determine which fractions contained properly folded scCl5, we pooled and concentrates the fractions indicated by the boxes on the chromatogram. The separate scCl5 peaks were used to coat fluorescent streptavidin beads and used to stain ML1 and K562 cells, as a control soluble γ9δ2TCR Cl5 beads were used. Only one peak, fractions 7-8, contained scCl5 that was able to stain K562 to similar levels as soluble Cl5 and did not stain ML1 (figure 1D). Although our observations implied that only a small part, 15-20%, of scCl5 is expressed as a properly folded protein that is able to bind recognized target cells, it did prove that γ9δ2TCRs can be expressed in single chain viable domains format, which is a prerequisite for using scTvs for affinity maturation with phage display.

**Figure 1.**
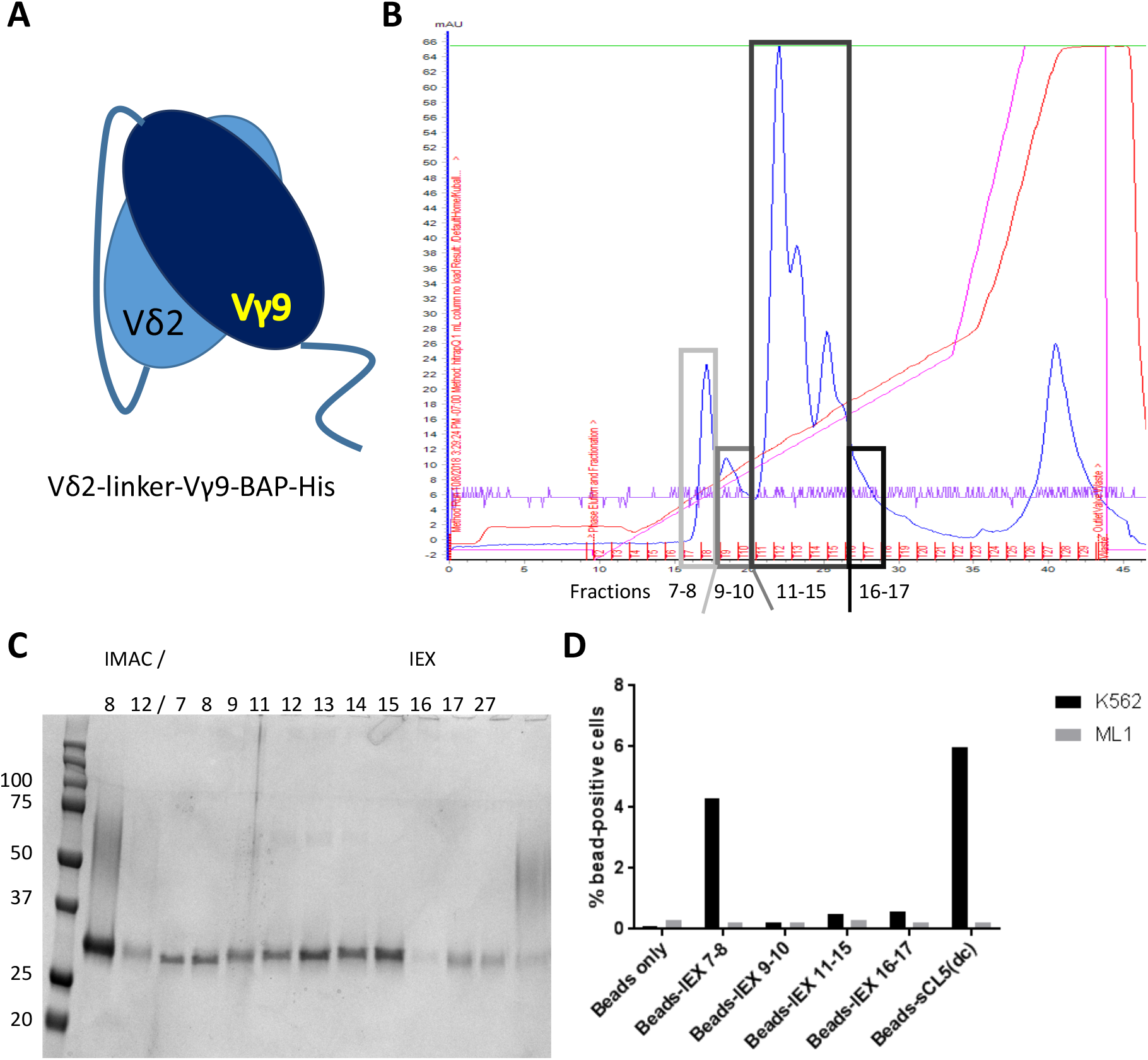
a fraction of the expressed single chain Cl5 binds to K562 cells. (A) schematic overview of the scCl5 construct, V2 domain is linked to the V9 domain using a flexible linker, attached to the C-terminus of the V9 domain are a biotin acceptor peptide (BAP) and a poly-histidine tag. (B) chromatogram of the anion exchange chromatography step of scCl5. Indicated fractions were analyzed by SDS-PAGE (C), pooled and tested for their ability to stain ML1 and K562 cells using fluoresent beads (D).

### Expression of scCl5 and the fusion protein scCl5-coat protein III in the periplasm of E.coli

After establishing that scCl5 was able to recognize K562 cells we cloned the scCl5 construct in the phagemid vector pUR8100, containing a pelB leader sequence for periplasmic localization and a Myc-His tag for detection. After the His-tag an amber stop codon is located followed by coat-protein III (gIII), in *E.coli* strains that can suppress the amber stop codon, this codon is translated 10-50% of the time in a glutamine leading to a scCl5-gIII fusion protein (figure 2A). In the presence of a helper phage this coat protein containing scCl5 will be incorporated in the phage particle.

**Figure 2.**
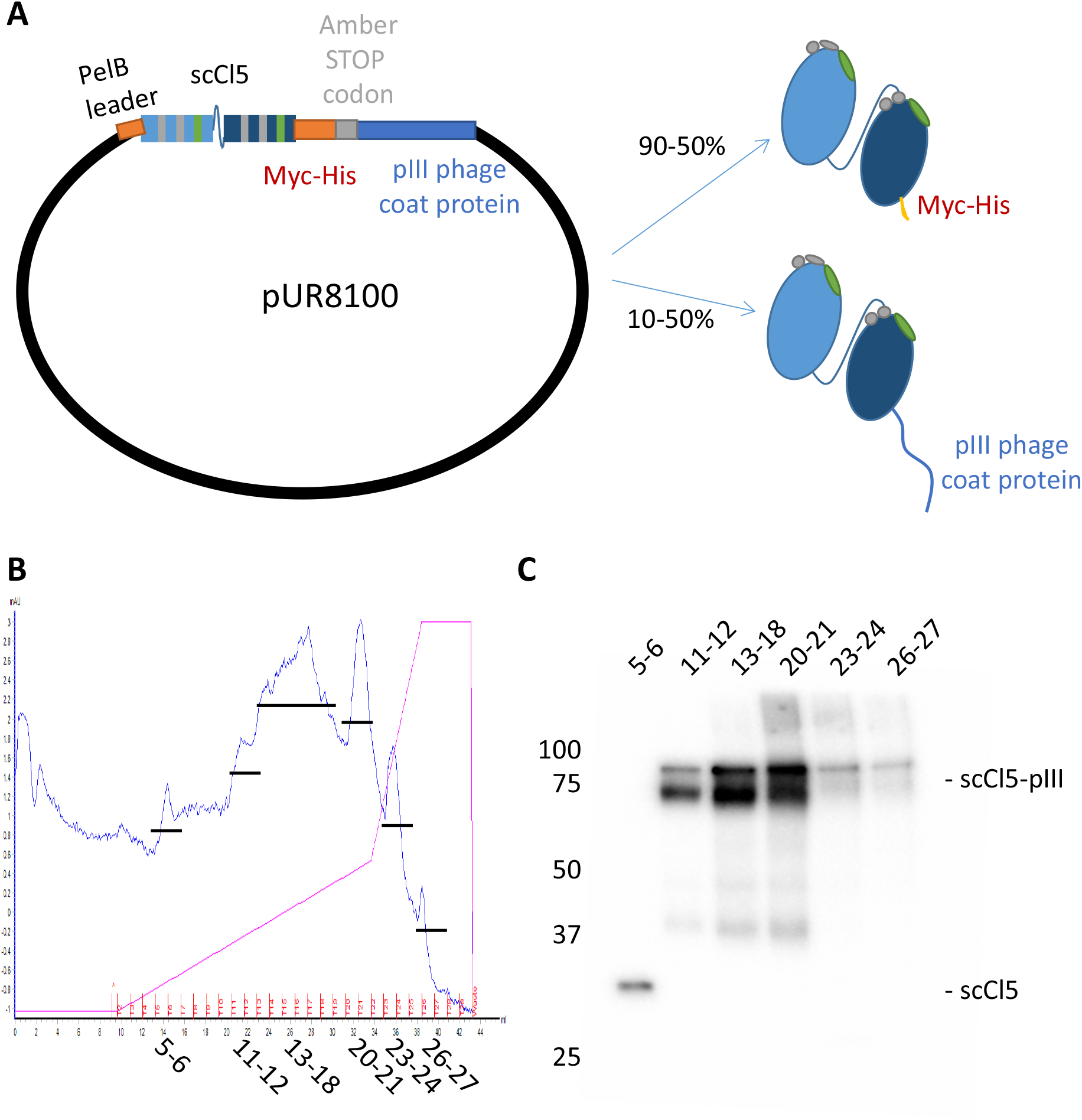
Expression of scCl5 in *E. coli* JM109. (A) schematic overview of bacterial expression plasmid pUR8100 containing scCl5. 5’ of the scCl5 gene is the PelB leader sequence and 3’the gene is linked to a Myc-His tag followed by a Amber stop codon and the pIII phage coat protein. The Amber stop codon can be suppressed by certain *E.coli* strains and results in the translation of the fusion protein at 10-50% efficiency. (B) chromatogram of the anion exchange chromatography step of scCl5/scCl5-pIII. (C) Indicated fractions of the IEX were pooled and analyzed by Western blot using an anti-polyHistidine antibody for detection.

We used the scCl5 containing phagemid to transform *E.coli* JM109, a strain capable of amber suppression, for large scale, simultaneous, expression of scCl5 and scCl5-gIII fusion proteins at 37°C for 3 hours. The periplasmic content was extracted using freeze thaw protocol in TBS, this extract was loaded on a nickel column to enrich for his-tagged proteins. The eluted fractions containing scCl5 and scCl5-gIII were further purified using anion exchange, as for the expression in HEK293F cells, many different peaks were observed in the chromatogram (figure 2B). As the peaks did not contain a lot op protein, judged by the low absorbance at 280 nm, the content of the different peaks was analyzed by Western blot (figure 2C). The first peak, fractions 5-6, contained scCl5, while all other peaks contained the scCl5-gIII fusion protein.

From this exercise we could draw several conclusions. First scCl5 is poorly expressed in the periplasm of *E.coli*, which might be due to poor stability or folding potential, as in mammalian cells also only 15-20% of the secreted scCl5 was functional. Second, the expression of scCl5-gIII fusion proteins is much better, compared to scCl5 alone, even though this fusion protein has a lower chance to be made due to the amber stop codon. It’s not uncommon that fusion proteins could rescue the expression of poorly folded proteins (14,15), but it does not always result in functional protein. Lastly, most of the fractions containing the scCl5-gIII protein contain two bands, indicating that the processing/folding was incomplete, as the expected size of the fusion protein should be around 75 kDa.

### Design and preparation of the D12G12 library

Supported by the positive results of scCl5-bead staining of K562 and the periplasmic expression of scCl5-gIII fusion protein, we started with the design of the library for affinity maturation of γ9δ2TCR. The phospho-antigen reactivity of the γ9δ2T cell subset is mediated by very diverse repertoire of CDR3δ and CDR3γ sequences of different lengths. In our design we will use the CDR3 sequences of Cl5, as this is the most potent TCR we have to date. We decided to introduce variation in the CDR1 and CDR2 of both the γ and the δ chain, in order to get a highly diverse library that could enhance binding next to CDR3 mediated interactions. Based on the surface accessibility of the CDR residues of G115, we decided to introduce the mutations on the following positions, for CDR1γ 7 amino acids at IMGT positions Gly27-Thr37, for CDR2γ 3 amino acids at IMGT positions Tyr58-Gly63, for CDR1δ 6 amino acids at IMGT positions Glu28-Tyr37, and for CDR2δ 3 amino acids at IMGT positions Glu56-Asp65. Each amino acid position was covered by an equimolar mix of codons encoding 19 amino acids, cysteine was excluded to prevent undesired disulfide bond formation. Combining the mutations in these four CDRs cumulated to a theoretical diversity of ~2*10^24 of the “D12G12” library (figure 3A).

**Figure 3.**
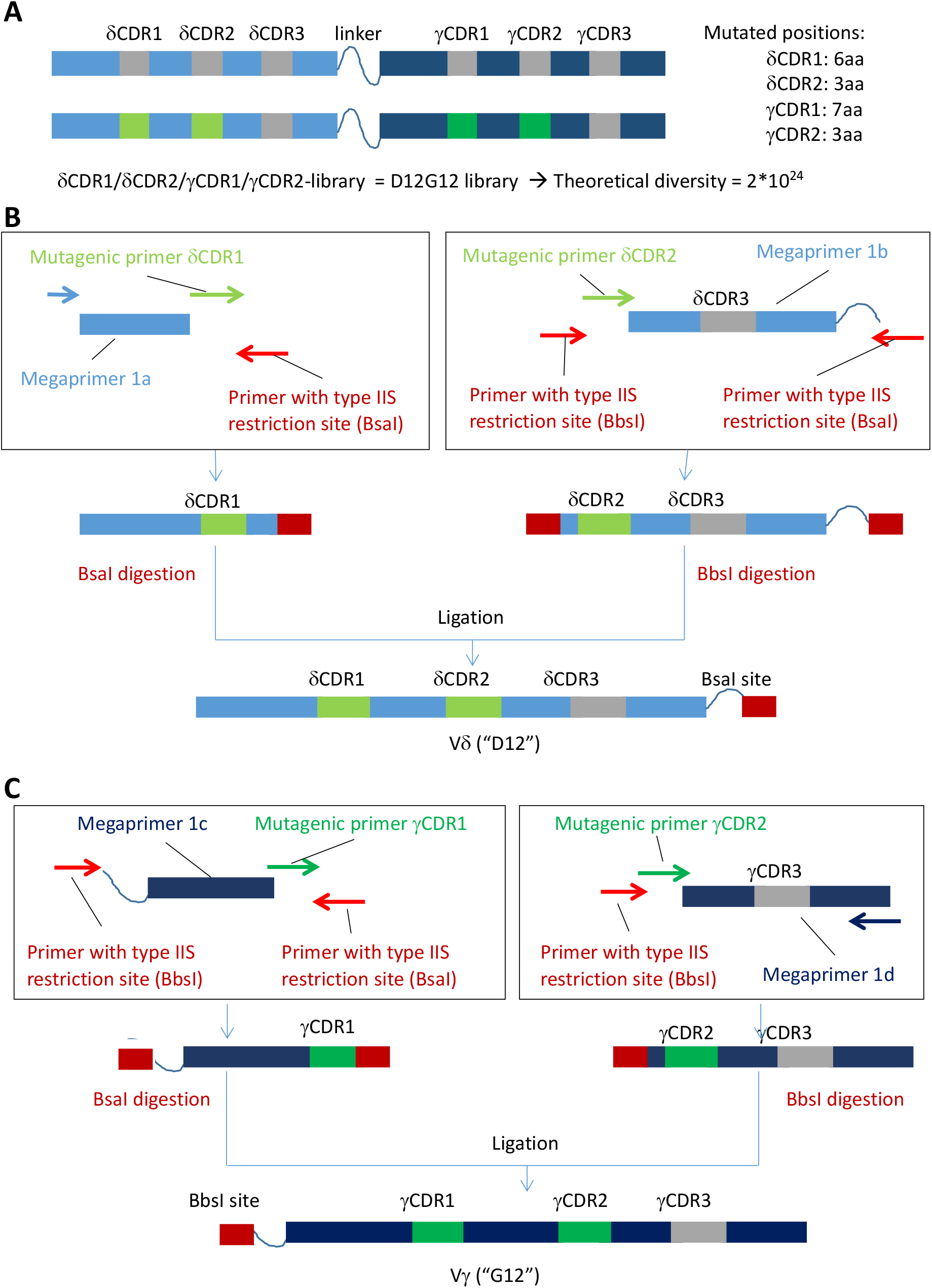
Design of the scCl5 CDR1&2 library. (A) The germline encoded CDR1 and CDR2 nucleotide sequences of Vδ2 and Vγ9 are replaced by 111 codons, an equimolar mix of trinucleotides coding for 19 amino acids except cysteine. Schematic overview of the assembly of the (B) Vδ2 “D12” and (C) Vγ9 “G12” library in multiple PCR and ligation steps.

The mutagenic primers were introduced in the scCl5 DNA by multiple PCRs, first four megaprimers were created by standard PCR, which served as scaffolds for the introduction of the mutagenic primers. All four mutagenic primers were fused to the megaprimers in four separate reactions (figure 3B and C), resulting in single PCR products (figure 4A). These products were flanked by BsaI and/or BsbI Type IIS restriction sites, that allow for seamless cloning of the different PCR products (figure 3B and C). In practice the ligation efficiency was low, around 20-30% of the input, CDR1δ + CDR2δ or CDR1γ + CDR2γ, was ligated to form the “D12” or “G12” genes (figure 4A). The final step ligation step of “D12” to “G12” to obtain “D12G12” again had an efficiency of 20%. This ligation inefficiency resulted in a theoretical drop in library diversity to around 10^22, but in practice, due to the limited amount of DNA in the reactions, this diversity was much lower, approximately 10^11. To be able to use the “D12G12” library for phage display it has to be ligated in the pUR8100 phagemid vector, which reduced the final diversity to around 10^6.

**Figure 4.**
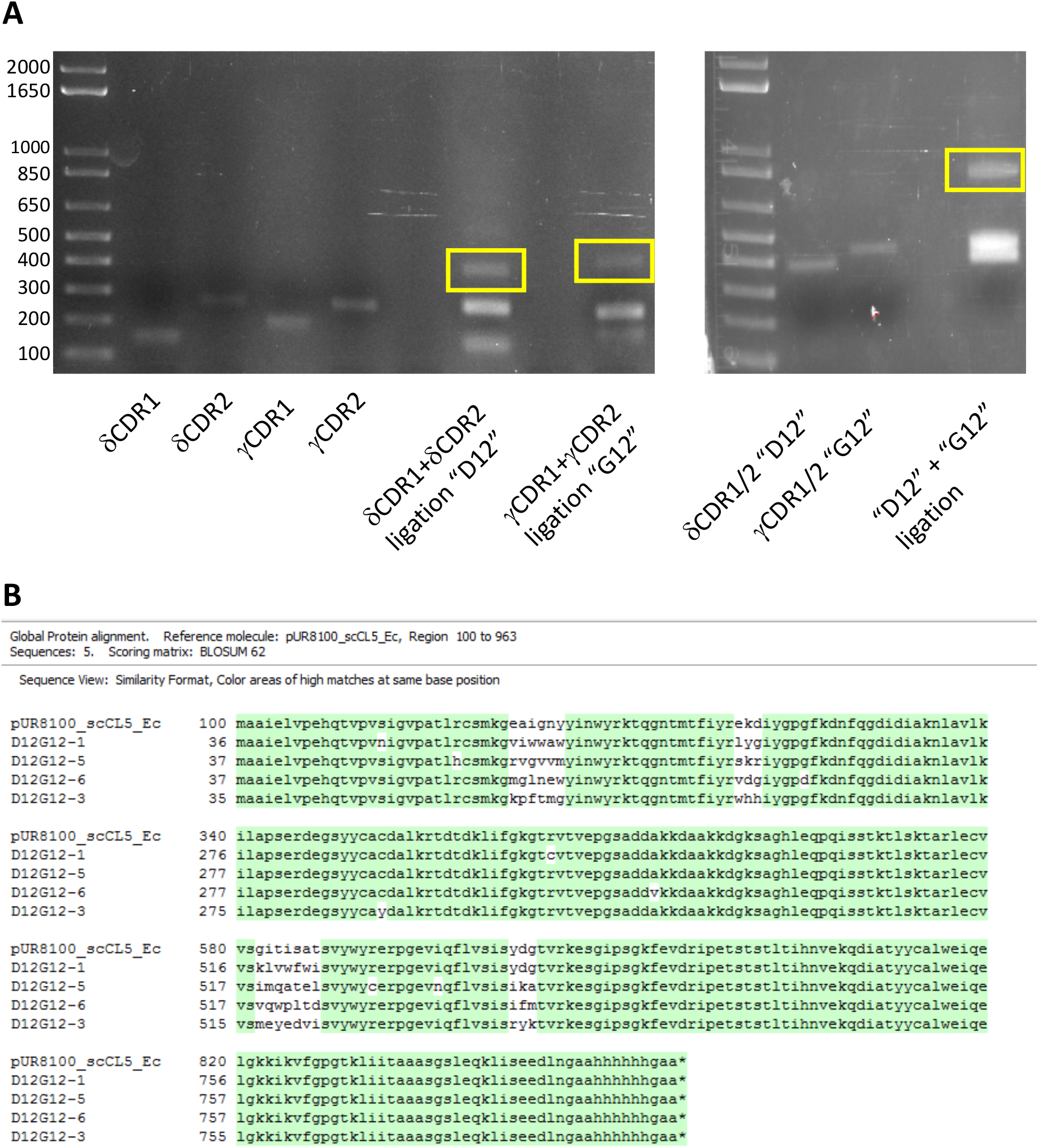
Preparation and quality control of the scCl5 “D12G12” library. (A) Analysis PCR and ligation products on a agarose gel. Ligation products of the correct size, indicated by the yellow box, were used for further cloning. (B) Sequence alignment of 4 randomly picked clones from the scCl5 “D12G12” library against the scCl5 sequence. Sequence highlighted in green corresponds to the scCl5 sequence. Alignment was made in Clone Manager Professional 9.

To assess the quality of the library we randomly selected 6 clones for sequencing. Two of the sequences contained mutations outside the CDRs leading to premature stop codons or frameshift mutations. The other four sequences contained some missense mutations outside the CDR1s and CDR2s, but could be translated to scTCR. Most importantly, all mutated positions contained unique amino acids, indicating that the practical diversity of 10^6 would not be influenced much by the library quality (figure 4B).

### Selection of scCl5 “D12G12” clones interacting with pamidronate treated K562 cells

Our strategy to obtain affinity matured scCl5 clones consisted of multiple bio-panning steps. The phage particles expressing the “D12G12” library were first incubated to the non-recognized ML1 cell line, which should reduce the number of “sticky” phages and library clones with altered specificity. The unbound fraction was incubated with pamidronate treated K562 cells and these cells were washed extensively. The bound phages were eluted and used for a second round of bio-panning and for a screen to identify strong binders (figure 5A).

**Figure 5.**
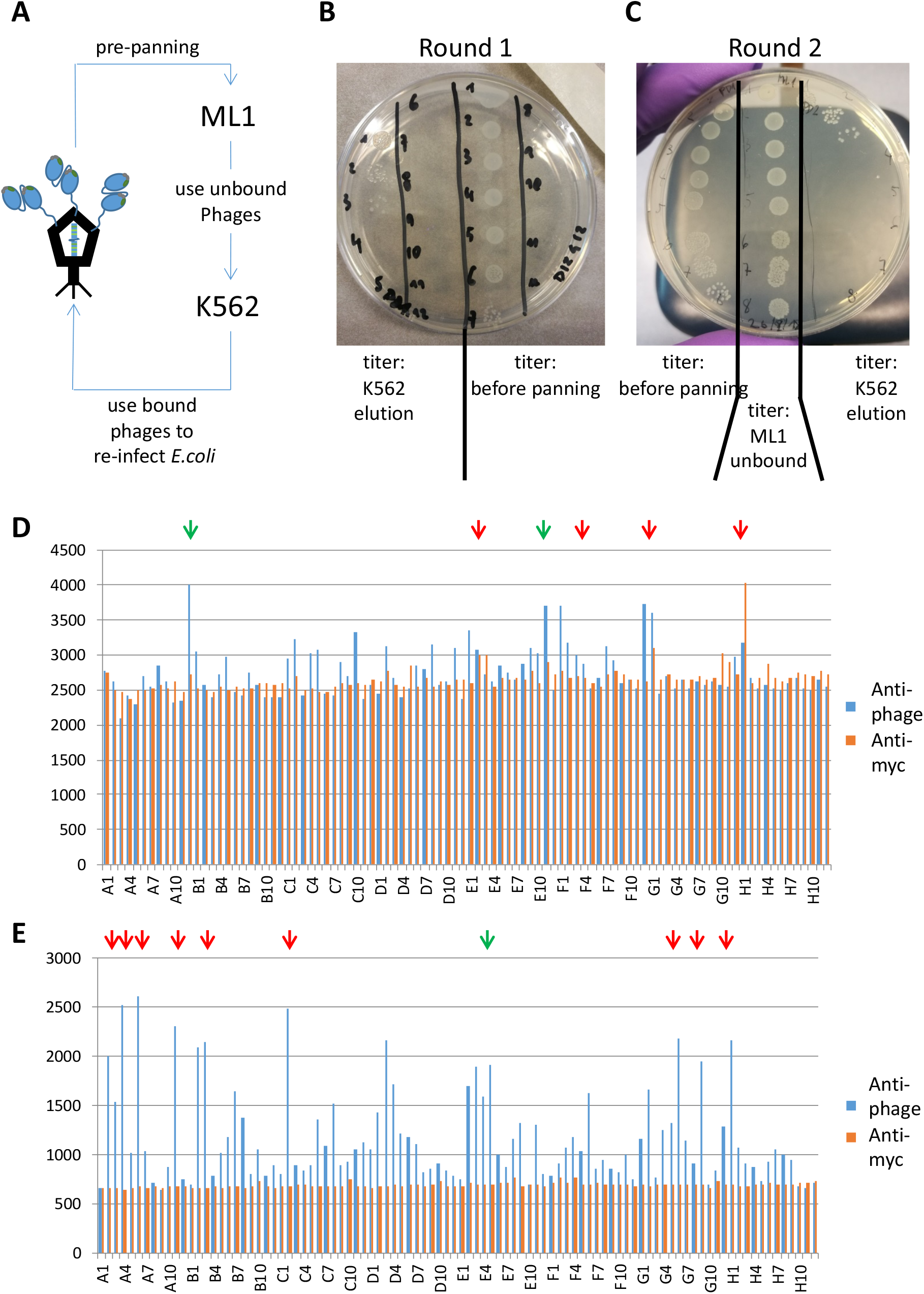
Phage display of scCl5 “D12G12” library. (A) Bio-panning strategy for the enrichment of scCl5-library phages with enhanced affinity for pamidronate treated K562 cells. Phage titers were determines using serial dilutions of phages infected *E.coli* using indicated phages fractions of the first round of enrichment (B) and the second round of enrichment (C). 94-phage clones of the K562-bound fraction, of the (D) first and (E) second panning round, were expressed as scTCR or phage particle and binding of these scTCR and phage particles to K562 was assessed using FACS. Phage particles were detected using a biotinylated anti-M13 (phage) antibody in combination with streptavidin PE and the scCl5-mutants were detected using a biotinylated anti-Myc antibody in combination with streptavidin-PE. Green arrows above are clones that have been sequenced and don’t have any nonsense/frameshift mutations, while red arrows indicate clones with these mutations (see supplemental figure 1).

In the first round of bio-panning we started with a phage titer of >10^11 pfu/ml, calculated by a serial dilution of re-infected *E.coli* (figure 5B), which means that each clone of the “D12G12” library is present ~10^5 times. After the pre-panning on ML1 cells the phage titer reduced to ~10^8 pfu/ml, indicating that a part of the phages was lost due to non relevant interaction. There was a further reduction when we determined the titer of the K562 eluted phages, ~10^6 pfu/ml, which indicates some form of selection (figure 5B).

The phagemid vectors of the eluted phages were amplified by reinfection on *E.coli* TG1 and a new phage preparation was made using the M13KO3 helper phage. Again the starting phage titer was high >10^11 pfu/ml, but this time there was no reduction in the phage titer after the pre-panning step on ML1 cells (figure 5C). This suggested that most “D12G12” library clones that interact with the non-recognizes cell line ML1 were lost in the first round of bio-panning, which points to specific enrichment of the library. This time we observed a further reduction in phage titer of the K562 eluted phages, indicating a further selection of the library (figure 5C).

The observed reduction in phage titer after both rounds of bio-panning was very encouraging, however to obtain specific clones, they have to be assessed on individual level for specific binding to K562. We randomly selected 94 clones of round 1 and 94 clones of round 2 and induced the expression of scCl5-mutant and scCl5mutant-gIII as done above. The periplasmic extracts were used to stain pamidronate treated K562 cells. Bound scCl5-mutants were detected using an anti-Myc tag antibody and the phages were detected using an anti-M13-phage antibody. Hardly any round 1 eluted clones showed increased binding to K562 as scTCR, which might be due to the poor expression we observed before. As scTCR-gIII fusion protein we could identify multiple clones with increased binding to K562 (figure 5D). The number of scTCR-gIII fusion protein clones from the second round of bio-panning that had increased binding was much greater than after round one (figure 5E). Again, the scTCR format did not show any binding. The larger number of K562 binding scTCR-gIII fusion proteins after round 2 compared to round 1 suggests that there was indeed an enrichment of K562-specific phages. To determine the sequences of these clones, 6 clones of round 1 and 10 clones of round 2 were send for sequencing. Unfortunately, many of the clones were incorrect, indicated by a red arrow (figures 5D & E), most incorrect clones had frameshift mutations, premature stop codons or the sequence couldn’t be aligned to scCl5 (supplemental figure 1A). In the end we were only able to identify three clones of these two rounds of phage display that could be translated in scTCR and one additional clone from an independent round of phage display with the same library (supplemental figure 1B). We conclude, that we have selected for random clones, of which many did not contain a correct scTCR sequence.

### Exploring alternatives for successful phage display

Although scFv and scTv fragments are most commonly used for phage display, other formats have been successfully used as well. Constructs comprising both the variable and constant domains of αβTCRs were incorporated in phage coat proteins and were used to enhance the affinity of several αβTCRs to the desired MHC-peptide complexes (16,17).

From our previous experience with soluble γδTCRs we know that they can be expressed in mammalian cells, however we never tested the expression of soluble γδTCRs in *E.coli*. To this end we ordered codon optimized constructs for the γ9δ2TCR Cl5 and the γ4δ5TCR C132 which contained an introduced cysteine in the constant domains for interchain disulfide bond formation, analogous to the interchain disulfide bond used in αβTCRs (12).

These constructs were ligated in 2 different plasmids, pMEK219 for periplasmic expression of soluble double chain γδTCR (dcTCR) and in phagemid pUR8100 for the expression of γδTCR-gIII fusion protein (figure 6A).

**Figure 6.**
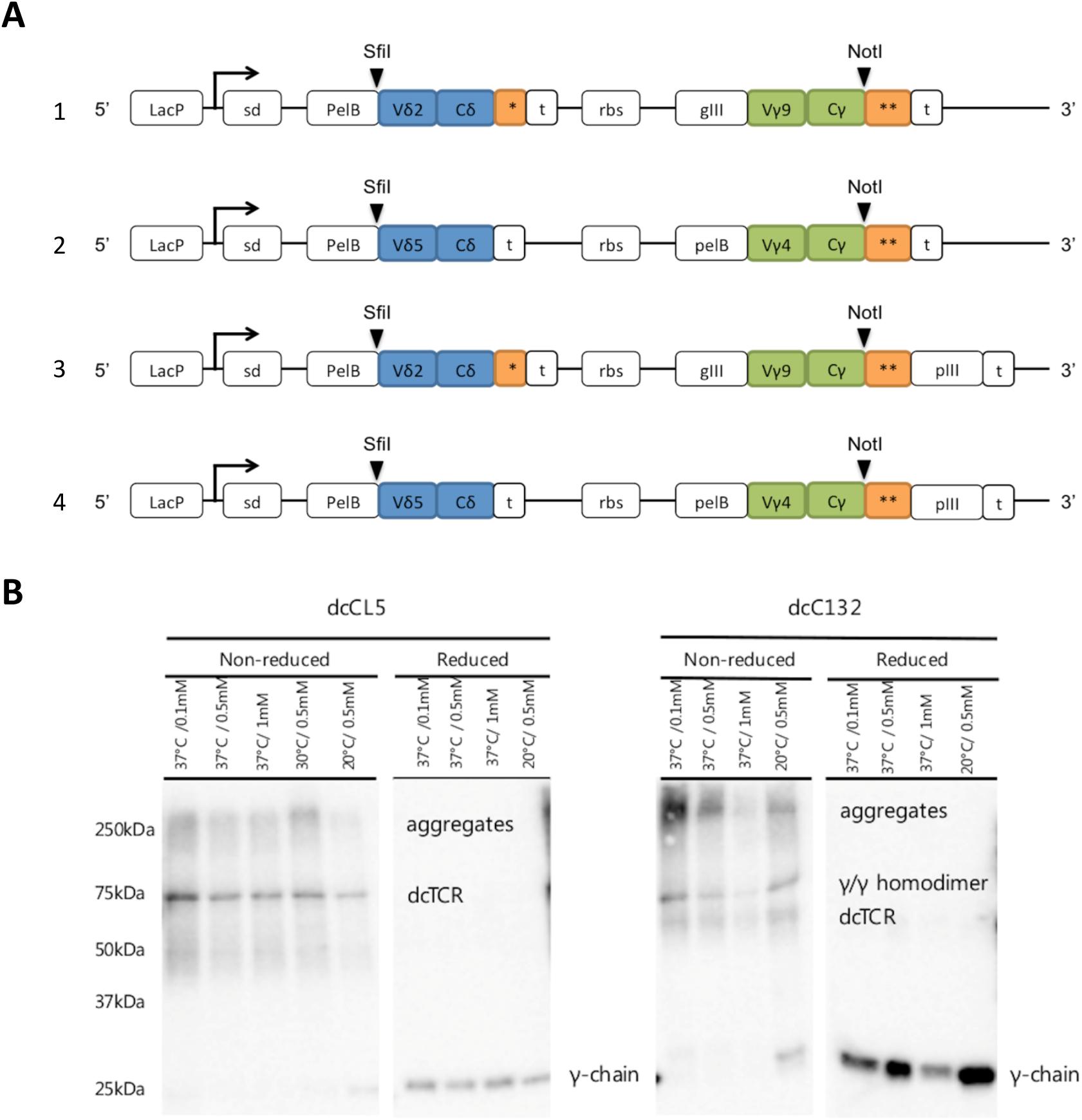
Double chain γδTCR expression in *E.coli* periplasm. (A) Schematic maps of the dcTCR encoding vector constructs (1) dcCL5 in pMEK219, (2) dcC132 in pMEK219, (3) dcCL5 in pUR8100 and (4) dcC132 in pUR8100. Abbreviations: lacP:lac promoter, sd: Shine-Dalagarno sequence, t: T7 terminator, rbs: ribosome binding site. SfiI, NotI: restriction enzyme sites. *Biotin Acceptor Peptide tag. ** c-Myc and His6-tags. (B) analysis of the periplasmic expression of the dcCL5 and dcC132 clones in pMEK219 in XL1-Blue under different expression conditions (temperature and [IPTG]). The samples have been blotted under non-reduced and reduced (addition of dithiothreitol) conditions. The his-tag, attached to the γ-chain, is detected via a primary anti-his antibody and a secondary anti-mouse antibody conjugated to HRP.

We assessed the periplasmic expression of both dcTCRs under a variety of expression conditions. Expression yields of proteins in *E.coli* can be optimized by reducing the temperature, which slows down the metabolism in the cells and thus the protein production rate, which might reduce misfolding of proteins. Another way to reduce the protein production rate is to use less IPTG, which lowers the number of copies of the gene of interest that are transcribed. For both dcCl5 as dcC132 reducing the IPTG concentration to 0.1 mM seemed to increase the formation of disulfide bonded TCR dimer compared to 0.5 and 1 mM slightly (figure 6B). However, under reducing conditions these differences were not that obvious. Moreover, under non-reducing conditions there was a smear visible in all lanes and for dcC132 there was an additional band around 75 kDa, this indicated poorly folded protein and the formation of γ-γ homodimers, respectively, as no additional bands are observed under reducing conditions (figure 6B).

Additional attempts to scale up the expression for the purification of dcTCR expressed in the periplasm of *E.coli* were unsuccessful. Due to the poor of expression the soluble dcTCRs, no further attempts were made to test the expression of the gIII-fusion protein for phage display.

### Stabilization of scTCR constructs

Both the scTCR as the dcTCR formats poorly express as soluble proteins in the periplasm of *E.coli*. Many others have encountered this scFv and scTv and their solution was to stabilize the single chain fragments by screening for stabilizing mutants (18,19). As read out they used binding to their desired antigen, which for γ9δ2TCR is not well defined.

As our aim is to identify expression enhancing mutations, we tested an experimental setup where we could detect periplasmic expression levels of his-tagged protein in a 96-wells format. *E.coli*, transformed with periplasmic expression constructs for a nanobody (Nb), scCl5, and scC132, were grown in 96-wells deep well blocks and protein expression was induced using 0.5 M IPTG for 3 h at 37C. Periplasmic extracts were loaded on nitrocellulose membrane using a dot-blot apparatus and his-tagged protein was detected using anti-polyhisitidine antibody. The three independent experiments showed mixed results, the well expressed Nb could be detected in every experiment, but the intensity between replicates varied (figure 7A). For the scTCRs, scCl5 and scC132, the detection was more variable in one experiment the proteins were hardly detected, while in the other two experiments some replicates showed detectable spots (figure 7A).

**Figure 7:**
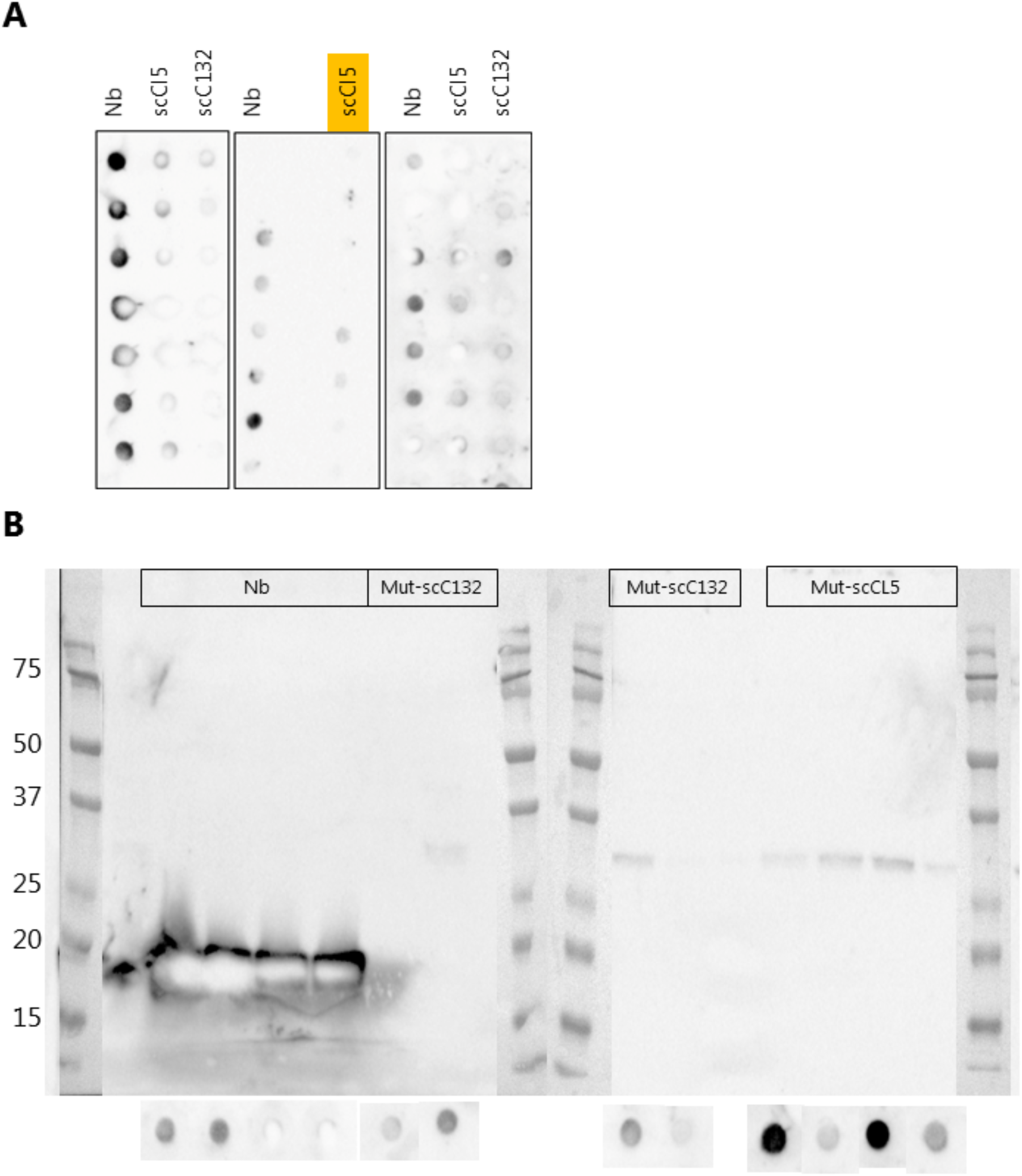
Screen to identify scTCR mutations increasing expression in *E.coli* (A) Analysis of periplasmic expression levels of a nanobody (Nb), scCI5, and scCl32 analyzed by Dot-Blot Periplasmic extracts were blotted on a nitrocellulose membrane and expressed recombinant protein was detected using a poly-histidine antibody (primary) and goat-anti-mouse-HRP (secondary). (B) Side-by-side analysis of periplasmic extracts of *E.coli* expressing Nb or scTCR-mutants by conventional Western blot and by Dot-Blot. Detection was done as in (A).

To determine how predictive this detection method was, we again expressed the Nb as positive control and some colonies containing mutants of scCl5 and scC132, that were created by random mutagenesis. Again, the periplasmic extracts were analyzed by dotblot and selected samples were analyzed by conventional Western blot to determine how these two analysis methods compared. For the Nb replicates there was very intense signal on Western blot, while the scTCR mutants showed only faint bands. The same samples on dot-blot did not had similar intensity, the Nb samples had much lower intensity compared to the scTCR samples (figure 7B).

Although the dot-blot analysis might have potential in the future for high throughput screening of scTCR expression in periplasmic extracts, the protocol needs to be further optimized to get representative results.

## Conclusion

Affinity maturation of γ9δ2TCR by using phage display was not successful. The largest hurdle was the periplasmic expression of γ9δ2TCR constructs in *E.coli* which is a prerequisite for successful phage display. The underlying reason for this lack of expression was the instability of the single chain (sc)TCR format. Expression of scTCR formats in HEK293F cells yielded only 15-20% correctly folded scTCR.

---

**Supplemental figure 1.**
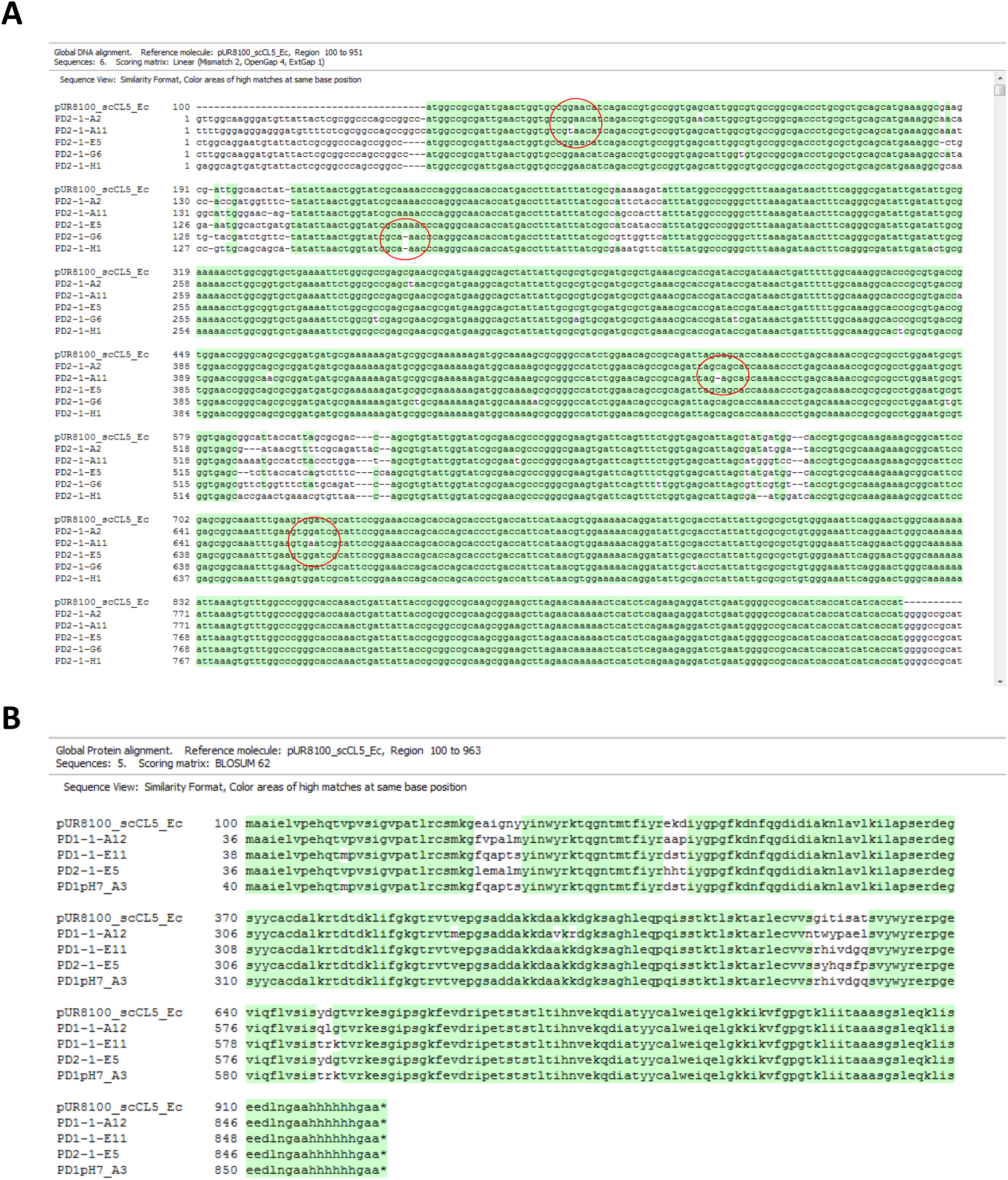
Sequences of phage clones in the K562 bound fraction. (A) Sequencing results of 5 clones with elevated phage binding to K562 after the second round of panning (Figure 5 E). Red circles indicate nonsense or frameshift mutations. (B) Alingment of amino acid sequences of 4 scCl5-phage clones without any nonsense or frameshift mutations. All alignments were made in Clone Manager Professional 9.

## Appendix 1: Sequences

### Expression constructs

Nucleotides in **Bold** were ordered at BaseClear the remainder of the sequences were plasmid encoded.

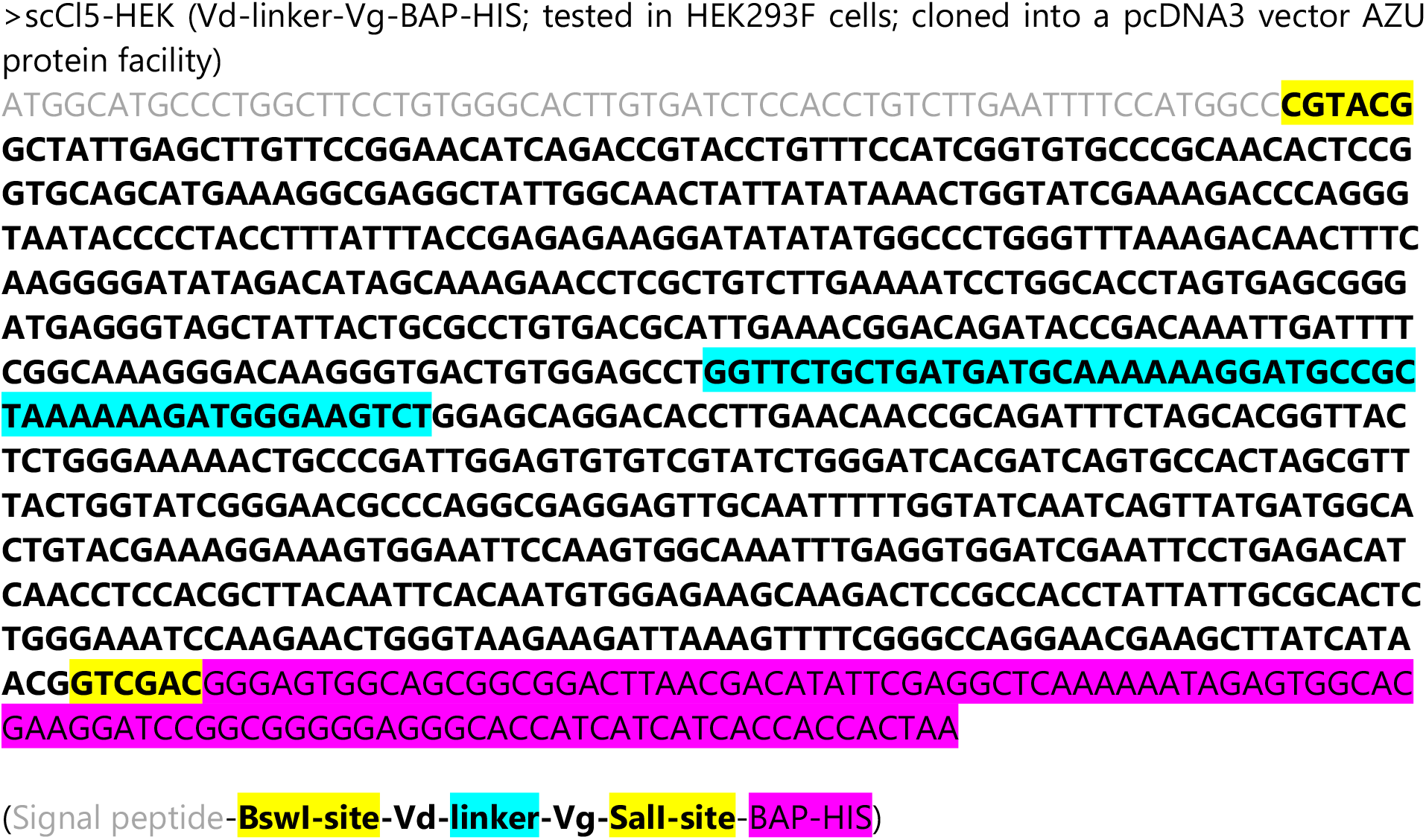

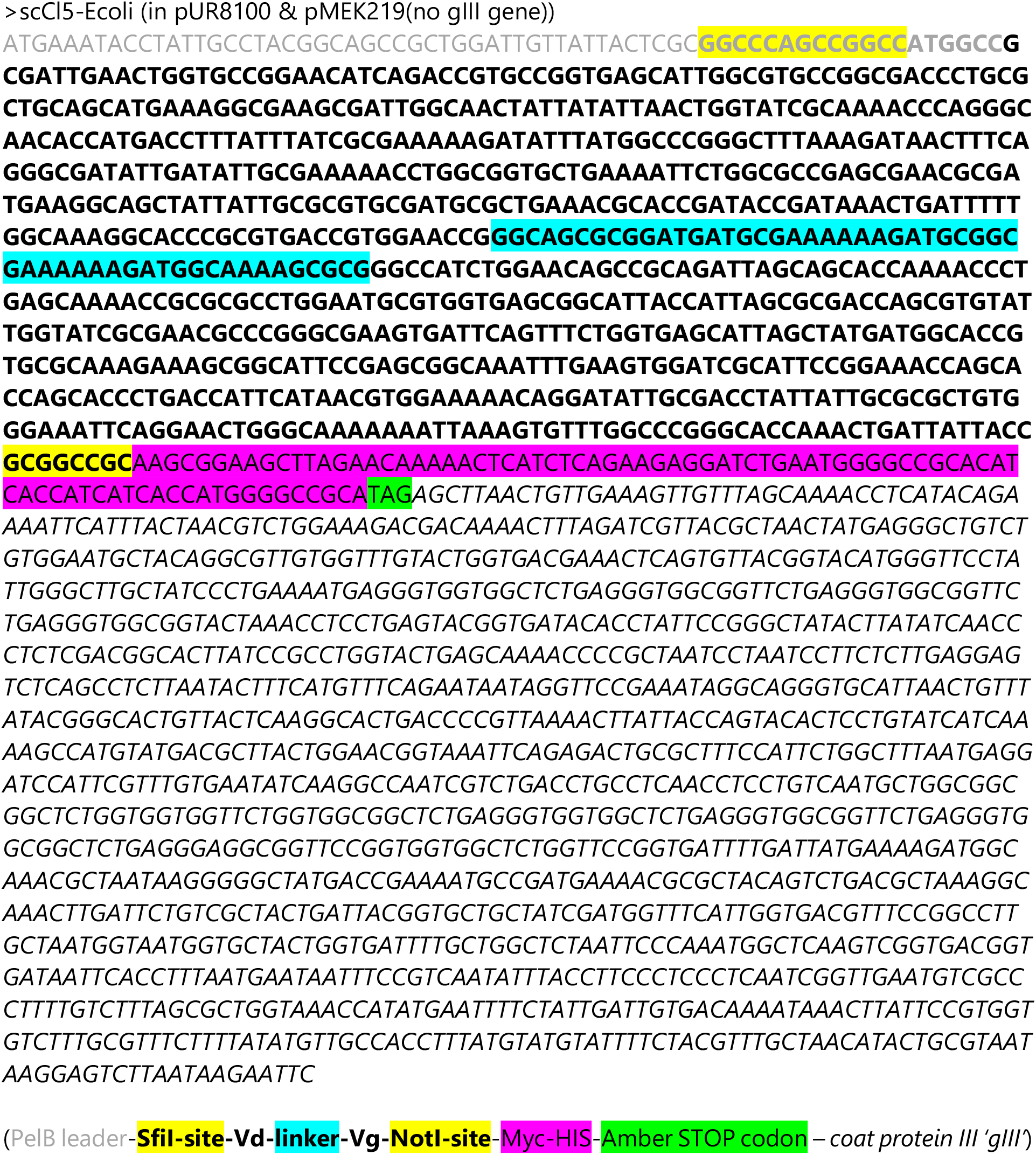

**Supplemental Table 1.**
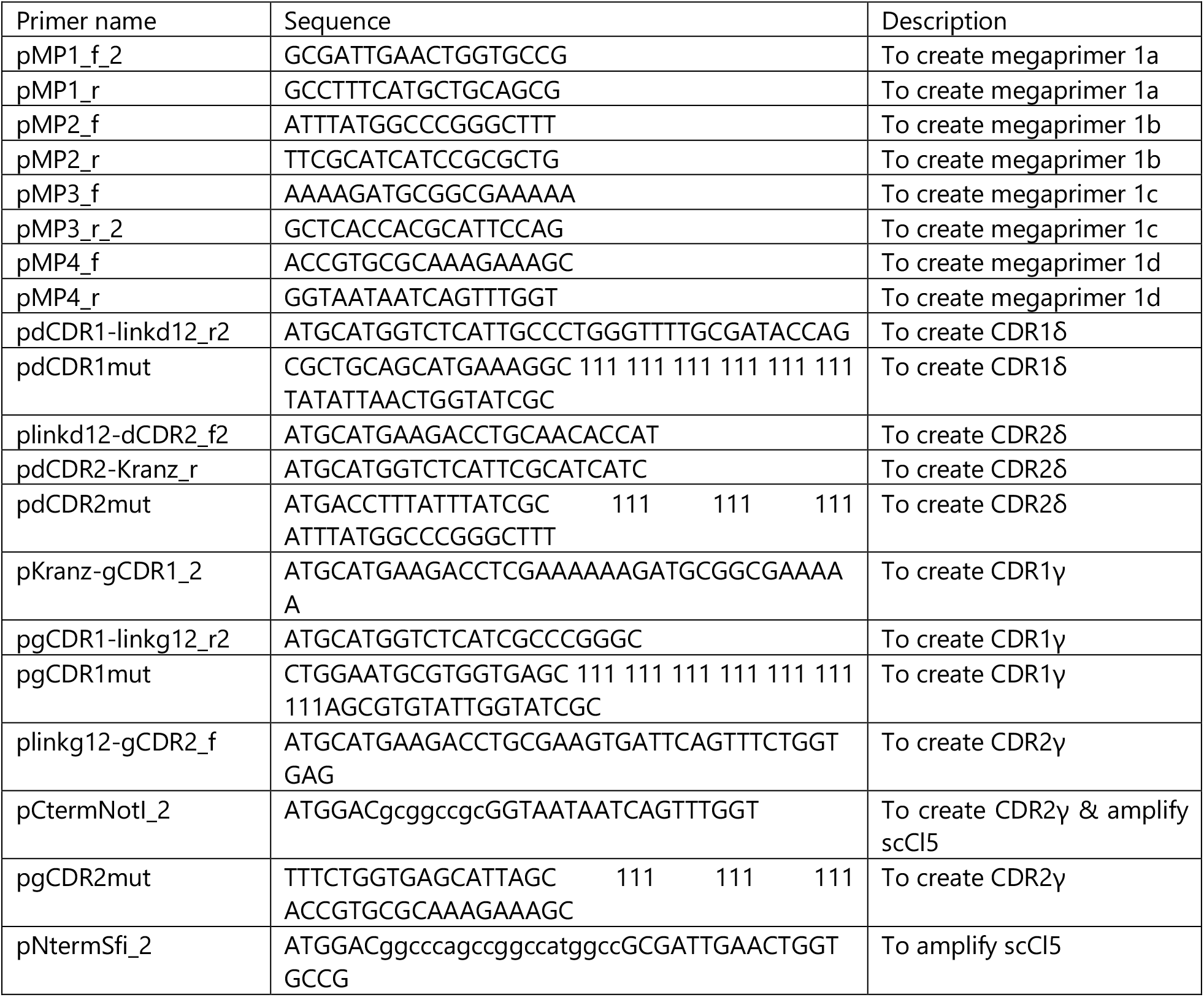
Overview of primers used to create scCl5”D12G12” library. 111 = equimolar mix of trinucleotides coding for 19 aa (no cysteine)

**Supplemental Table 2.**
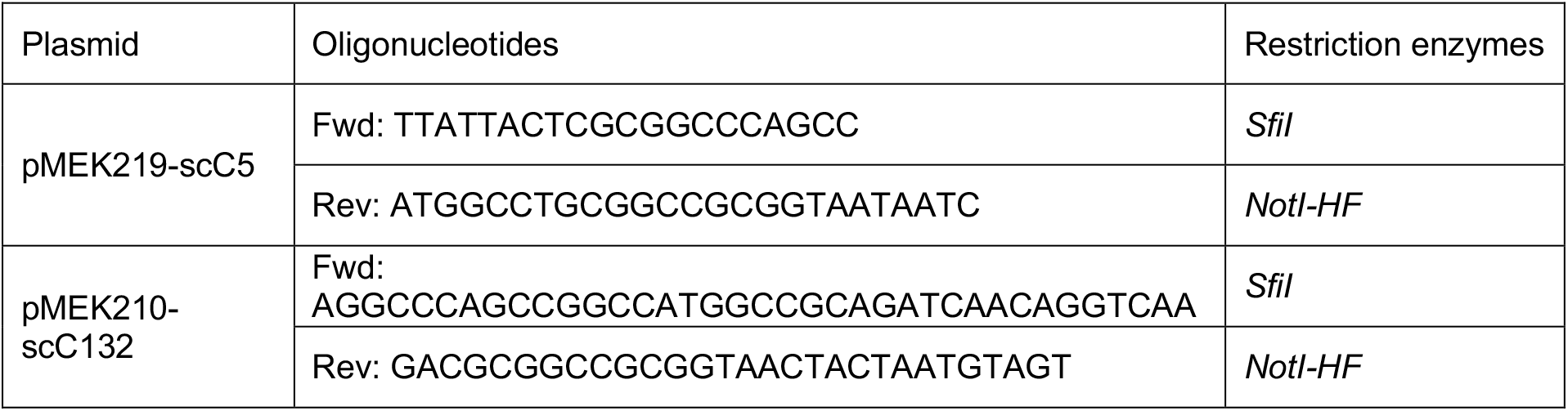
Primer sequences used for random mutagenesis of scCl5 and scC132.

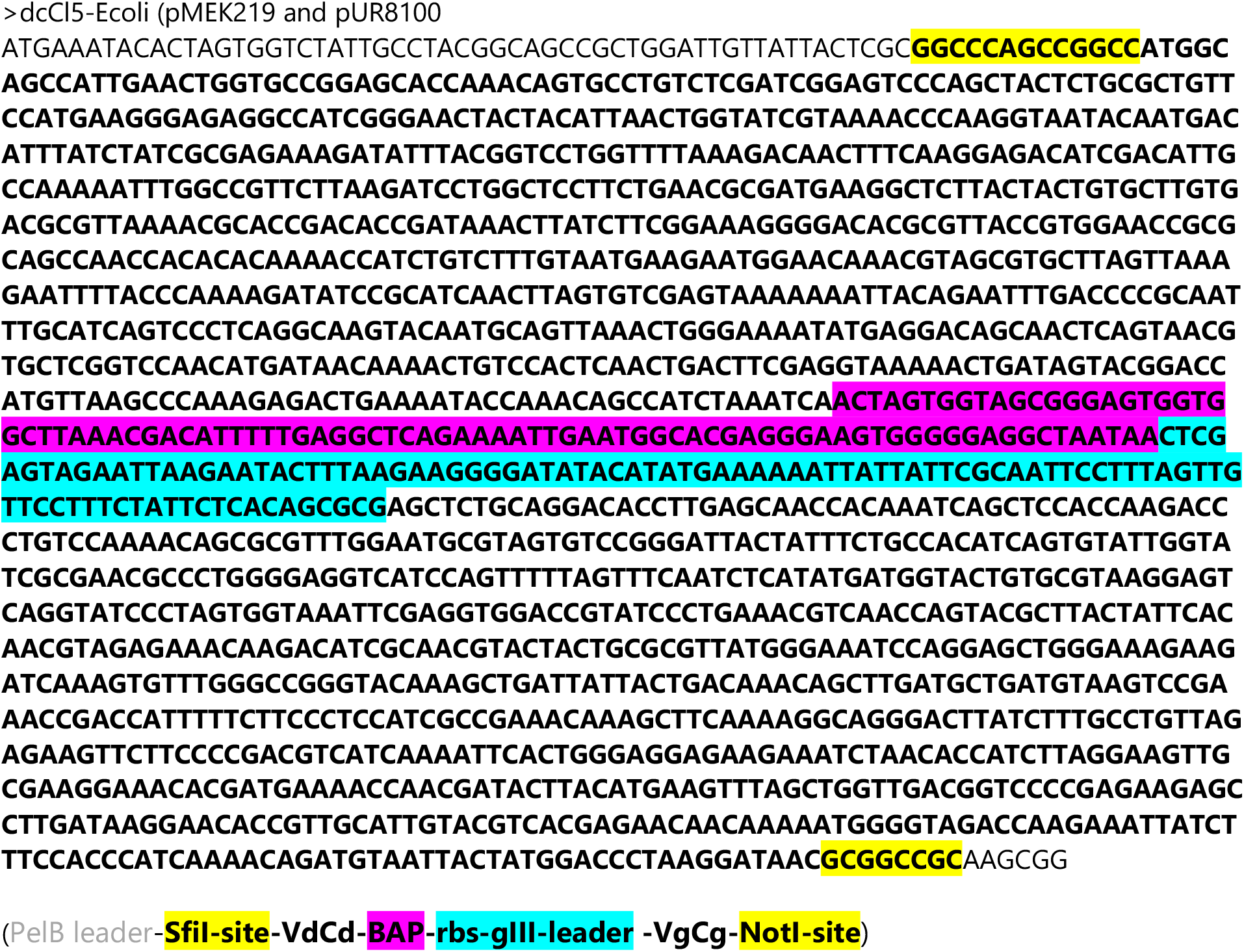

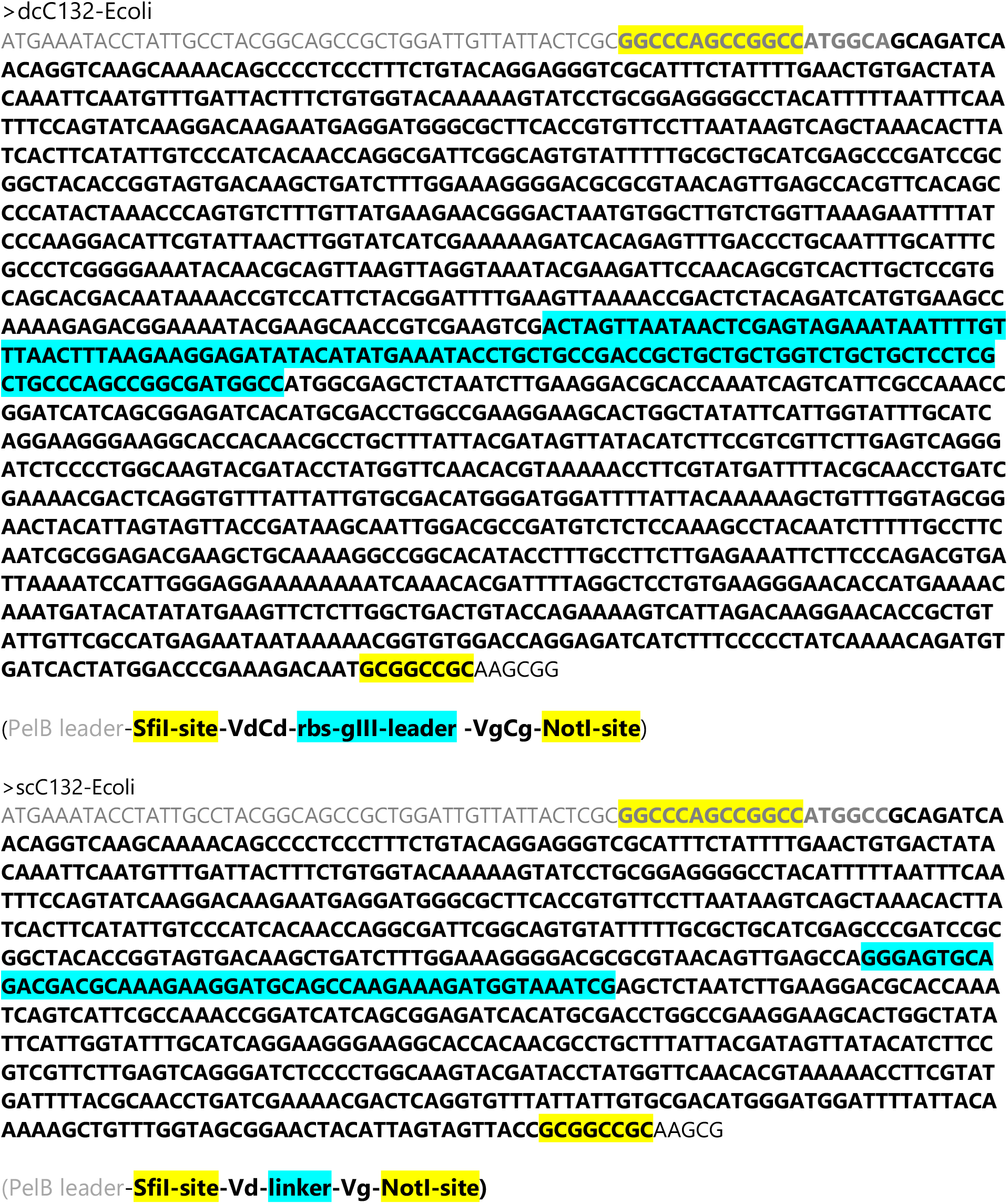

### Clones obtained by phage display

>PD1-1-A12

GTGGCAATGATGTATTACTCGCGGCCCAGCCGGCCATGGCCGCGATTGAACTGGTGCCGGAACATCA GACCGTGCCGGTGAGCATTGGCGTGCCGGCGACCCTGCGCTGCAGCATGAAAGGCTTCGTTCCAGCA CTGATGTATATTAACTGGTATCGCAAAACCCAGGGCAACACCATGACCTTTATTTATCGCGCAGCACCA ATTTATGGTCCGGGCTTTAAAGATAACTTTCAGGGCGATATTGATATTGCGAAAAACCTGGCGGTGCTG AAAATTCTGGCGCCGAGCGAACGCGATGAAGGCAGTTATTATTGCGCGTGCGATGCGCTGAAACGCA CCGATACCGATAAACTGATTTTTGGCAAAGGCACCCGCGTGACCATGGAACCGGGCAGCGCGGATGA TGCGAAAAAAGATGCGGTGAAAAGAGATGGCAAAAGCGCGGGCCATCTGGAACAGCCGCAGATTAG CAGCACCAAAACTCTGAGCAAAACCGCGCGCCTGGAATGCGTGGTGAACACCTGGTACCCAGCAGAA CTGAGCGTGTATTGGTATCGCGAACGCCCGGGCGAAGTGATTCAGTTTCTGGTGAGCATTAGCCAGCT GGGTACCGTGCGCAAAGAAAGCGGCATTCCGAGCGGCAAATTTGAAGTGGATCGCATTCCGGAAACC AGCACCAGCACCCTGACCATTCATAACGTGGAAAAACAGGATATTGCGACCTATTATTGCGCGCTGTG GGAAATTCAGGAACTGGGCAAAAAAATTAAAGTGTTTGGCCCGGGCACCAAACTGATTATTACCGCG GCCGCAAGCGGAAGCTTAGAACAAAAACTCATCTCAGAAGAGGATCTGAATGGGGCCGCACATCACC ATCATCACCATGGGGCCGCATAGAGCTTAACTGTTGAAAGTTGTTTAGCAAAACCTCATACAGAAAATT CATTTACTAACGTCTGGAAAGACGACAAAACTTTAGATCGTTACGCTAACTATGAGGGCTGTCTGTGGA ATGCTACAGGCGTTGTGGTTTGTACTGGTGACGAAACTCAGTGTTACGGTACATGGGTTCCTATTGGGC TTGCTATCCCTGAAAAATGAAGGTGGTGGCTCTGAAGGGTGGCGGTTCTGAAGGTTGCCGGTTCTGAA GGGTGGCGGTACTAAACCTCCCTGAATACGGGGATACCACCTATTCCGGGCTAAACTTATATCAACCC TCTCGAAGGGACTTTATCCGCCTGGGACTGAACAAAACCCGCTAATCCAAAACCTTCTTCTTGGAGGA ATTCAACCCCTTAAAAATTCCAGTTTTCCAAAAATAAGGTTCCCAAAAAAGAACGGGGGGCTTAACTGT TTTATCGGGCCCTGGTTTCAAGGGGCTGACCCCGTTAAAACTATTACCAAAACTCCCGTTTATCAAAAA CCGTGTGGAGCCTCTTGGGGACGGAATTCAAAAAGCGTTTCTTTTCTGCCTTTAGAAGAAACCTTTTTT GAATAAAAGGAAACCAATACCCTCACCTCCTTATGTCGGGGGCGCTGGGTGTGGTTGGGTGTCTTAAG AGTGGACTGAAAATCCCCAGTGGGTGGTTTGTGCGTGGGCGTTCGCCCCGCGCCTGTCCTGTTTACAA TTATATCCCGTTTTAAAATTAAAAAAAGAGAAAAATCCCTCTCTCCTAGATCCGTGTTCCGTTTCGCGTC CTCTTCCTTCCCTCTACCTCCTCTTTCTTTTCATTTTGCCTTTATCCCTAATATTATAAACCTGATATCTTTA GCGTAAGTTTCACGGGCGCGGACCCTTAACGGCACGCCGAATTTCACGTGGTGGGAAAATAAAGCGA

>PD1-1-E11

GTGGCAAGGGATGTTATTACTCGCGGCCCAGCCGGCCATGGCCGCGATTGAACTGGTGCCGGAACAT CAGACCATGCCGGTGAGCATTGGCGTGCCGGCGACCCTGCGCTGCAGCATGAAAGGCTTCCAGGCAC CAACCTCTTATATTAACTGGTATCGCAAAACCCAGGGCAACACCATGACCTTTATTTATCGCGATTCTA CCATTTATGGCCCGGGCTTTAAAGATAACTTTCAGGGCGATATTGATATTGCGAAAAACCTGGCGGTG CTGAAAATTCTGGCGCCGAGCGAACGCGATGAAGGCAGCTATTATTGCGCGTGCGATGCGCTGAAAC GCACCGATACCGATAAACTGATTTTTGGCAAAGGCACCCGCGTGACCGTGGAACCGGGCAGCGCGGA TGATGCGAAAAAAGATGCGGCGAAAAAAGATGGCAAAAGCGCGGGCCATCTGGAACAGCCGCAGAT TAGCAGCACCAAAACCCTGAGCAAAACCGCGCGCCTGGAATGCGTGGTGAGCCGTCATATCGTTGAT GGTCAGAGCGTGTATTGGTATCGCGAACGCCCGGGCGAAGTGATTCAGTTTCTGGTGAGCATTAGCAC CCGTAAAACCGTGCGCAAAGAAAGCGGCATTCCGAGCGGCAAATTTGAAGTGGATCGCATTCCGGAA ACCAGCACCAGCACCCTGACCATTCATAACGTGGAAAAACAGGATATTGCGACCTATTATTGCGCGCT GTGGGAAATTCAGGAACTGGGCAAAAAAATTAAAGTGTTTGGCCCGGGCACCAAACTGATTATTACCG CGGCCGCAAGCGGAAGCTTAGAACAAAAACTCATCTCAGAAGAGGATCTGAATGGGGCCGCACATCA CCATCATCACCATGGGGCCGCATAGAGCTTAACTGTTGAAAGTTGTTTAGCAAAACCTCATACAGAAA ATTCATTTACTAACGTCTGGAAAGACGACAAAACTTTAGATCGTTACGCTAACTATGAGGGCTGTCTGT GGAATGCTACAGGCGTTGTGGTTTGTACTGGTGACGAAACTCAGTGTTACGGTACATGGGTTCCTATTG GGCTTGCTATCCCTGAAAATGAGGGTGGTGGCTCTGAAGGTGGCGGTTCTGAAGGTGGCGGTTCTGAA GGTGGCGGTACTAAACCTCCTGAATACGGTGATACACCTATTCCGGGCTATACTTATATCAACCCTCTC GACGGCCTTATCCGCCTGGTACTGAACAAAACCCGCTAAACCAATCCCTTCCTTGAGGAATCCAACCC CTTAAAACTTCCTGTTTCCAAAAATAAGGTTCCAAAAAAGCCGGGGGCTTTAACGTTTTAACGGGCCTG TTACCCAGGCCTGAACCCGTTAAACTATTACAAAAACTCCGGTTTCCAAAACCGTGTGAGCCCTCTGG GAACGGAATACAAAACGCTCTCTTTCTTGCTTTATGGAAACATTTTTTGGAATAAAGGAATCTCTACGC CCACCTCCAATTGGGGGGGCGGCGGTGTGGGTGTTTTTCCCCCGAGGAGAACAATAAAGGCGAGAGG ATAAGGGAAGCGGAAAGGGAATAGGGGCTCCCCCCCTCCTATAAAATAAAAAAAAAAAATTAAAAAA AAAAAAAAAGGCCCCCCAAATTCTGTGAACATGTTAAAAGAGGTAGGGGGTACGTGAAATAAGGATA AAATACATATTTAAATCTTTACTAAATCGATTAAAGATATCAAAATCCAATTCACAGACACAATGGCTTT TCGTTTTTACATTTTGGGTTTTGGAACCGG

>PD2-1-E5

CTGGCAGGAATGTATTACTCGCGGCCCAGCCGGCCATGGCCGCGATTGAACTGGTGCCGGAACATCA GACCGTGCCGGTGAGCATTGGCGTGCCGGCGACCCTGCGCTGCAGCATGAAAGGCCTGGAAATGGCA CTGATGTATATTAACTGGTATCGCAAAACCCAGGGCAACACCATGACCTTTATTTATCGCCATCATACC ATTTATGGCCCGGGCTTTAAAGATAACTTTCAGGGCGATATTGATATTGCGAAAAACCTGGCGGTGCT GAAAATTCTGGCGCCGAGCGAACGCGATGAAGGCAGCTATTATTGCGCGTGCGATGCGCTGAAACGC ACCGATACCGATAAACTGATTTTTGGCAAAGGCACCCGCGTGACCGTGGAACCGGGCAGCGCGGATG ATGCGAAAAAAGATGCGGCGAAAAAAGATGGCAAAAGCGCGGGCCATCTGGAACAGCCGCAGATTA GCAGCACCAAAACCCTGAGCAAAACCGCGCGCCTGGAATGCGTGGTGAGCTCTTACCATCAGTCTTTC CCAAGCGTGTATTGGTATCGCGAACGCCCGGGCGAAGTGATTCAGTTTCTGGTGAGCATTAGCTATGA TGGCACCGTGCGCAAAGAAAGCGGCATTCCGAGCGGCAAATTTGAAGTGGATCGCATTCCGGAAACC AGCACCAGCACCCTGACCATTCATAACGTGGAAAAACAGGATATTGCGACCTATTATTGCGCGCTGTG GGAAATTCAGGAACTGGGCAAAAAAATTAAAGTGTTTGGCCCGGGCACCAAACTGATTATTACCGCG GCCGCAAGCGGAAGCTTAGAACAAAAACTCATCTCAGAAGAGGATCTGAATGGGGCCGCACATCACC ATCATCACCATGGGGCCGCATAGAGCTTAACTGTTGAAAGTTGTTTAGCAAAACCTCATACAGAAAATT CATTTACTAACGTCTGGAAAGACGACAAAACTTTAGATCGTTACGCTAACTATGAGGGCTGTCTGTGGA ATGCTACAGGCGTTGTGGTTTGTACTGGTGACGAAACTCAGTGTTACGGTACATGGGTTCCTATTGGGC TTGCTATCCCTGAAAATGAAGGTGGTGGCTCTGAGGTTGGCGGTTCTGAGGGTGGCGGTTCTGAAGGT GGCGGTACTAAACCTCCTGAATACGGTGATACACCTATTCCGGGCTATACTTATATCAACCCTCTCCAA CGGCACTTATCCGCCTGGAACTGAGCAAAACCCCGCTAATCCTAATCCCTTTCTCTGGAGAGATCCCA GCCTCTTAAAACTTTCCATGTTTCAGAAAAATAAGGTTCCCAAATAAGGCAGGGGTGCTATAACTGGTT TATACGGGGCACTGGTTAATCAAGGCCCTGAACCCCGTTAAAACTTTTTTCCCGAACCCTCCTGGTATC TCAAAAAACCTGTGTTGAACTTTTCTTGGAACGGGAATTTCAAAACCC

>PD1pH7_A3

AGTTGAGAATTCATGTTATTACTCGCGGCCCAGCCGGCCATGGCCGCGATTGAACTGGTGCCGGAACA TCAGACCATGCCGGTGAGCATTGGCGTGCCGGCGACCCTGCGCTGCAGCATGAAAGGCTTCCAGGCA CCAACCTCTTATATTAACTGGTATCGCAAAACCCAGGGCAACACCATGACCTTTATTTATCGCGATTCT ACCATTTATGGCCCGGGCTTTAAAGATAACTTTCAGGGCGATATTGATATTGCGAAAAACCTGGCGGT GCTGAAAATTCTGGCGCCGAGCGAACGCGATGAAGGCAGCTATTATTGCGCGTGCGATGCGCTGAAA CGCACCGATACCGATAAACTGATTTTTGGCAAAGGCACCCGCGTGACCGTGGAACCGGGCAGCGCGG ATGATGCGAAAAAAGATGCGGCGAAAAAAGATGGCAAAAGCGCGGGCCATCTGGAACAGCCGCAGA TTAGCAGCACCAAAACCCTGAGCAAAACCGCGCGCCTGGAATGCGTGGTGAGCCGTCATATCGTTGAT GGTCAGAGCGTGTATTGGTATCGCGAACGCCCGGGCGAAGTGATTCAGTTTCTGGTGAGCATTAGCAC CCGTAAAACCGTGCGCAAAGAAAGCGGCATTCCGAGCGGCAAATTTGAAGTGGATCGCATTCCGGAA ACCAGCACCAGCACCCTGACCATTCATAACGTGGAAAAACAGGATATTGCGACCTATTATTGCGCGCT GTGGGAAATTCAGGAACTGGGCAAAAAAATTAAAGTGTTTGGCCCGGGCACCAAACTGATTATTACCG CGGCCGCAAGCGGAAGCTTAGAACAAAAACTCATCTCAGAAGAGGATCTGAATGGGGCCGCACATCA CCATCATCACCATGGGGCCGCATAGAGCTTAACTGTTGAAAGTTGTTTAGCAAAACCTCATACAGAAA ATTCATTTACTAACGTCTGGAAAGACGACAAAACTTTAGATCGTTACGCTAACTATGAGGGCTGTCTGT GGAATGCTACAGGCGTTGTGGTTTGTACTGGTGACGAAACTCAGTGTTACGGTACATGGGTTCCTATTG GGCTTGCTATCCCTGAAAATGAAAGGTGGTGGCTCTGAAGGGTGGCGGTTCTGAAGGTGGGCGGTTCT GAAGGTGGCGGTACTAAACCTCCTGAATACGGTGATACACCAATTCCGGGCTATACTTATATCAACCC TCTCAACGGACCTTATCCGCCTGGGAACTGAAAAAAACCCCGCTAATCCAAACCCTTCTCTGAGGAGA TTCAGCCCTCTAAAATTTCATGGTTCAAAATTAAGGTTCCAAAAAAGCAGGGGGCTTTACTGTTTTAAG GGGCAGGTTATCCAGGGTTGGCCCCTTATATT

## References

1. Vyborova A, Beringer DX, Fasci D, Karaiskaki F, van Diest E, Kramer L, et al. gamma9delta2T cell diversity and the receptor interface with tumor cells. The Journal of clinical investigation 2020 doi 10.1172/JCI132489.

2. Grunder C, van DS, Hol S, Drent E, Straetemans T, Heijhuurs S, et al. gamma9 and delta2CDR3 domains regulate functional avidity of T cells harboring gamma9delta2TCRs. Blood 2012;120(26):5153–62.

3. Straetemans T, Grunder C, Heijhuurs S, Hol S, Slaper-Cortenbach I, Bonig H, et al. Untouched GMP-Ready Purified Engineered Immune Cells to Treat Cancer. Clinical cancer research: an official journal of the American Association for Cancer Research 2015;21(17):3957–68 doi 10.1158/1078-0432.CCR-14-2860.

4. Liddy N, Bossi G, Adams KJ, Lissina A, Mahon TM, Hassan NJ, et al. Monoclonal TCR-redirected tumor cell killing. Nat Med 2012;18(6):980–7 doi 10.1038/nm.2764.

5. Rapoport AP, Stadtmauer EA, Binder-Scholl GK, Goloubeva O, Vogl DT, Lacey SF, et al. NY-ESO-1-specific TCR-engineered T cells mediate sustained antigen-specific antitumor effects in myeloma. Nat Med 2015;21(8):914–21 doi 10.1038/nm.3910.

6. Harly C, Guillaume Y, Nedellec S, Peigne CM, Monkkonen H, Monkkonen J, et al. Key implication of CD277/butyrophilin-3 (BTN3A) in cellular stress sensing by a major human gammadelta T-cell subset. Blood 2012;120(11):2269–79.

7. Karunakaran MM, Willcox CR, Salim M, Paletta D, Fichtner AS, Noll A, et al. Butyrophilin-2A1 Directly Binds Germline-Encoded Regions of the Vgamma9Vdelta2 TCR and Is Essential for Phosphoantigen Sensing. Immunity 2020;52(3):487–98 e6 doi 10.1016/j.immuni.2020.02.014.

8. Palakodeti A, Sandstrom A, Sundaresan L, Harly C, Nedellec S, Olive D, et al. The molecular basis for modulation of human Vgamma9Vdelta2 T cell responses by CD277/butyrophilin-3 (BTN3A)-specific antibodies. JBiolChem 2012;287(39):32780–90.

9. Rigau M, Ostrouska S, Fulford TS, Johnson DN, Woods K, Ruan Z, et al. Butyrophilin 2A1 is essential for phosphoantigen reactivity by gammadelta T cells. Science 2020;367(6478) doi 10.1126/science.aay5516.

10. Sebestyen Z, Scheper W, Vyborova A, Gu S, Rychnavska Z, Schiffler M, et al. RhoB Mediates Phosphoantigen Recognition by Vgamma9Vdelta2 T Cell Receptor. Cell Rep 2016;15(9):1973–85 doi 10.1016/j.celrep.2016.04.081.

11. Richman SA, Aggen DH, Dossett ML, Donermeyer DL, Allen PM, Greenberg PD, et al. Structural features of T cell receptor variable regions that enhance domain stability and enable expression as single-chain ValphaVbeta fragments. Mol Immunol 2009;46(5):902–16 doi 10.1016/j.molimm.2008.09.021.

12. Boulter JM, Glick M, Todorov PT, Baston E, Sami M, Rizkallah P, et al. Stable, soluble T-cell receptor molecules for crystallization and therapeutics. Protein Eng 2003;16(9):707–11 doi 10.1093/protein/gzg087.

13. Richman SA, Kranz DM, Stone JD. Biosensor detection systems: engineering stable, high-affinity bioreceptors by yeast surface display. Methods Mol Biol 2009;504:323–50 doi 10.1007/978-1-60327-569-9_19.

14. Costa S, Almeida A, Castro A, Domingues L. Fusion tags for protein solubility, purification and immunogenicity in Escherichia coli: the novel Fh8 system. Front Microbiol 2014; 5:63 doi 10.3389/fmicb.2014.00063.

15. Ki MR, Pack SP. Fusion tags to enhance heterologous protein expression. Appl Microbiol Biotechnol 2020;104(6):2411–25 doi 10.1007/s00253-020-10402-8.

16. Gunnarsen KS, Hoydahl LS, Neumann RS, Bjerregaard-Andersen K, Nilssen NR, Sollid LM, et al. Soluble T-cell receptor design influences functional yield in an E. coli chaperone-assisted expression system. PLoS One 2018;13(4):e0195868 doi 10.1371/journal.pone.0195868.

17. Li Y, Moysey R, Molloy PE, Vuidepot AL, Mahon T, Baston E, et al. Directed evolution of human T-cell receptors with picomolar affinities by phage display. Nat Biotechnol 2005;23(3):349–54 doi 10.1038/nbt1070.

18. Kieke MC, Shusta EV, Boder ET, Teyton L, Wittrup KD, Kranz DM. Selection of functional T cell receptor mutants from a yeast surface-display library. Proc Natl Acad Sci U S A 1999;96(10):5651–6 doi 10.1073/pnas.96.10.5651.

19. Shusta EV, Holler PD, Kieke MC, Kranz DM, Wittrup KD. Directed evolution of a stable scaffold for T-cell receptor engineering. Nat Biotechnol 2000;18(7):754–9 doi 10.1038/77325.

